# Plateau-gated one-shot plasticity supports continual recognition memory

**DOI:** 10.64898/2026.09.04.749460

**Authors:** Guanchun Li, Sandro Romani, Jeffrey C. Magee

## Abstract

Biological memory systems store single experiences while continuing to learn, but how one-shot plasticity limits interference with existing memories is unclear. Behavioral timescale synaptic plasticity (BTSP) rapidly modifies synapses active within seconds of a dendritic plateau. We isolate its plateau-triggered component in a model where plastic weights and a stable instructive pathway jointly determine whether a plateau occurs, closing a feedback loop between the synaptic state and the plastic event that modifies it. For unstructured inputs, instantiated by independent uniform signed patterns, the dynamics reduce exactly to pathway alignment, which determines the fidelity of the instructed representation, the synaptic-turnover rate and the mean rewrite interval. For structured inputs, instantiated by correlated bimodal Curie–Weiss patterns, the instructive pathway biases which component of input structure enters the plastic synaptic state. In a BTSP-inspired continual-recognition network, combined instructive and plastic drives determined the selected memory unit for each one-shot write, whereas a Hebbian control used the same plastic weights for credit assignment and memory storage. The BTSP-inspired network remained accurate at longer repeat lags than Hebbian controls, an advantage that grew with network size, with both architectures optimized independently at every repeat lag. A reduced theory predicted held-out accuracy, lag capacity and dynamics of memory-trace strength directly from optimized parameters. It showed why intermediate proximal and distal coupling was optimal: proximal plastic drive guided plateau generation toward selected memory units, slowing synaptic turnover but limiting new encoding, whereas distal instructive drive enhanced familiar responses but could also make novel inputs appear familiar. The memory-trace strength in the rate-and-depth-matched Hebbian control still decayed faster and showed less effective credit assignment than in the BTSP-inspired network. These results connect dendritic plateau physiology to continual memory and support partial separation of allocation from storage as a mechanism for limiting interference during continual learning.

## Introduction

Biological memory systems must combine rapid acquisition with resistance to interference from subsequent experience. Rapid one-trial storage has long been proposed as a core hippocampal computation^1^. A dendritic plateau during a single traversal can induce a place field in a hippocampal CA1 pyramidal neuron^2–4^, and human observers can subsequently recognize thousands of images, each presented only once ^5,6^. By contrast, many conventional artificial neural networks are trained through repeated, incremental parameter updates and are susceptible to catastrophic forgetting when experiences are presented sequentially^7–10^. Determining how neural systems combine rapid encoding with persistent storage during ongoing learning is therefore a central problem in both neuroscience and machine learning.

Continual recognition memory—the online classification of each item in an uninterrupted stream as novel or previously encountered—provides a stringent network-level assay of this problem. Each novel item must be encoded after one presentation while later inputs continue to modify the same finite memory substrate. Recognition of prior occurrence can likewise be established after limited exposure in animals^11^, and theoretical familiarity networks have characterized storage in this regime ^12–15^. Here we use continual recognition as a benchmark to ask whether a stable, anatomically motivated instructive pathway can direct credit assignment by determining the selected memory unit through plateau generation. This model-level selection is related to the broader biological problem of neuronal memory allocation, in which neurons compete for participation in a memory trace ^16,17^.

The discovery of BTSP provided a concrete cellular mechanism for rapid reorganization of hippocampal representations ^3,18^. In CA1 and CA3 pyramidal neurons, a regenerative dendritic plateau potential lasting hundreds of milliseconds acts as an instructive plasticity event, rapidly modifying synaptic inputs active within a window extending for several seconds around the plateau^3,19,20^. The direction and magnitude of these changes depend on initial synaptic strength and input-plateau timing, allowing both potentiation and depression ^19^. Distal entorhinal input contributes to plateau initiation and instructs the resulting changes in spatial representations ^21,22^. BTSP therefore couples a threshold-like dendritic event to rapid, coordinated synaptic modification across a seconds-long eligibility window. This places it within the broader family of behavioral-timescale eligibility-trace and three-factor learning rules ^23^. Here, the regenerative dendritic plateau is the specific instructive event, distinguishing the mechanism from conventional Hebbian rules based on temporally coincident pre- and postsynaptic activity ^24^.

Classical theories of synaptic memory have largely been developed around pattern-driven local transition rules. Attractor-network theory established collective retrieval and high-load behavior for distributed patterns ^25–27^. Sparse binary and low-activity models showed how coding level and synaptic architecture reshape associative capacity^28–31^. Palimpsest and bounded-synapse models characterized forgetting under continual, often one-step, synaptic changes ^32–38^. Metaplastic and complex-synapse models showed how hidden states can prolong memory and constrain achievable memory trade-offs ^39–41^. Some frameworks already permit one-shot, stochastic or state-dependent changes, and selective updating can improve memory of correlated patterns with bounded synapses^42^. The additional structure considered here is not one-shot plasticity itself but a shared plasticity-inducing event. Pattern-defined eligibility is filtered through a dendritic plateau that the current weights help recruit, and the plateau in turn gates coordinated modification of those same weights. Analyzing this loop therefore requires the geometry of allowed transitions and the statistics of waiting times between them, in addition to conventional signal-to-noise arguments. Related studies have examined BTSP-dependent place-field formation, BTSP-based content-addressable storage and specialized online memory rules^43–45^. To our knowledge, however, no framework has jointly connected a shared, self-looping plasticity-inducing event whose occurrence depends on the current plastic state and a relatively stable instructive pathway, the stationary transition and waiting-time statistics produced by that event, and quantitative continual-recognition performance.

Here we develop such a framework, in which the instructive and plastic pathways have partly distinct roles in credit assignment and memory storage. A relatively stable instructive pathway biases event occurrence and, in the network, credit assignment to a selected memory unit, whereas a separate plastic pathway carries the changing synaptic representation. We first isolate the plateau-triggered component of BTSP as a gated jump process. Each input pattern proposes a new plastic synaptic state, and the integrated dendritic drive determines event occurrence. For unstructured inputs instantiated by independent uniform signed patterns, the resulting dynamics collapse exactly onto a single measure of pathway alignment between the instructive and plastic pathways. This reduction shows how the effective plateau threshold and the balance of proximal and distal drive govern event occurrence, preservation of the instructed representation, microscopic synaptic turnover and the mean rewrite interval. For structured inputs instantiated by correlated bimodal Curie–Weiss patterns, the theory shows how the instructive pathway biases which component of input structure is incorporated into the plastic synaptic state.

We then carry this organization into a BTSP-inspired continual-recognition network that includes the cross-time depression of BTSP. On every presentation, the current plastic drive and a stable distal instructive drive jointly determine normalized plateau probability and hence which memory unit gets written, whereas in the Hebbian control, the plastic weights alone determine credit assignment and store the memory. With each architecture optimized independently at each repeat lag, the BTSP-inspired network remains accurate at longer repeat lags than the Hebbian control at every tested width and input dimension, with a capacity advantage that grows across the tested grid. A reduced theory predicts held-out accuracy and lag capacity for BTSP-inspired network, and the strength of a tagged model memory trace, from the optimized parameters alone, with nothing fitted to the simulations. The theory further explains why intermediate coupling is beneficial. When proximal plastic drive contributed to credit assignment, the selected memory unit was more likely to be one already responsive to the current pattern, slowing subsequent synaptic turnover but leaving less room for potentiation during new encoding. Distal instructive drive instead strengthened memory-unit responses to familiar patterns without changing the underlying synaptic engram, but could also make some novel patterns appear familiar. In a matched trace intervention, a rate-and-depth-matched Hebbian trace control used the BTSP-inspired network learning rates and cascade depth, with its remaining parameters re-optimized at its own capacity. It nonetheless showed faster decay of the memory-trace strength. Together, these results suggest that allocating credit assignment and storage to partly distinct pathways, combined with sparse one-shot writes to selected memory units, can reduce interference during continual learning.

## Results

### One-shot plasticity as a gated jump process

Two experimentally established properties of BTSP motivate our theoretical formulation (Fig. 1a). First, plasticity is gated by a regenerative, threshold-like dendritic plateau, with event occurrence depending on the strength and temporal conjunction of entorhinal and CA3 drive^2,21^. Second, a plateau can modify inputs active within a temporal window extending for several seconds on either side of the event; the direction of modification depends in part on initial synaptic strength, permitting both potentiation and depression^3,19,20^. We isolate these two event-triggered properties and do not treat long-term stability of synaptic weights as a defining assumption.

**Figure 1:**
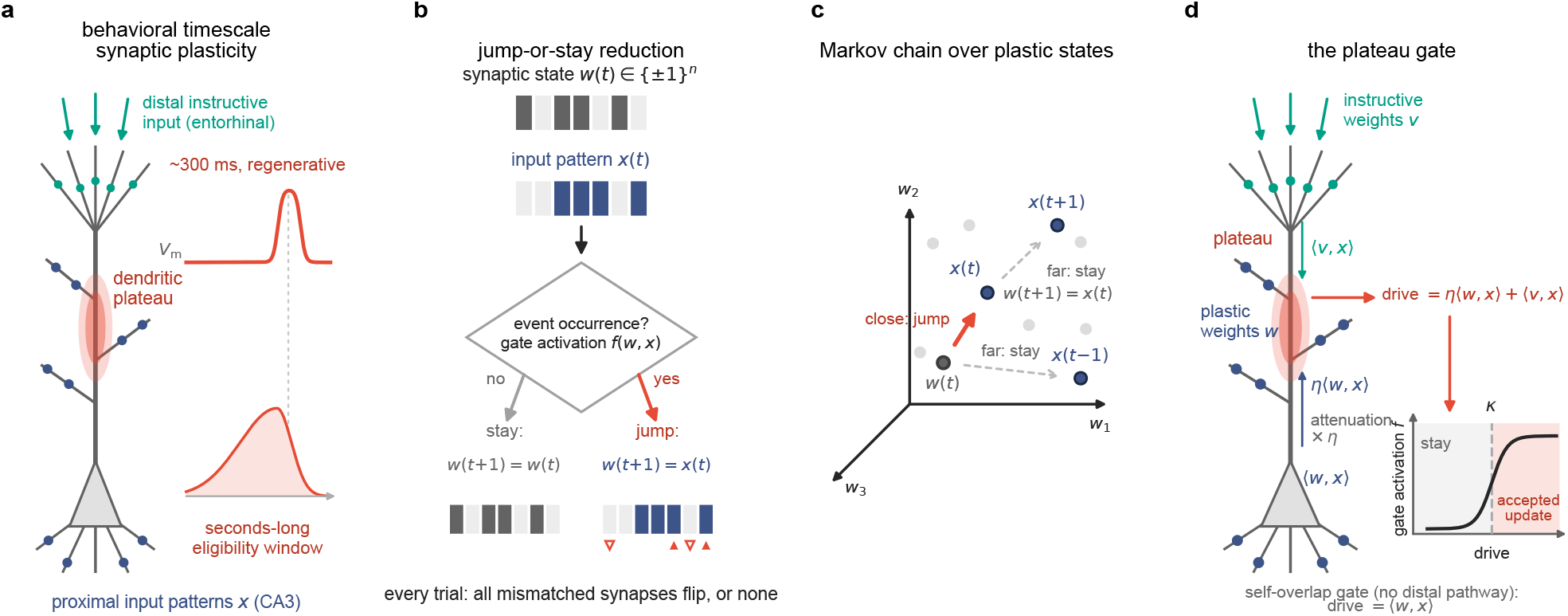
One-shot plasticity as a gated jump process. **a**, Behavioral timescale synaptic plasticity: a regenerative dendritic plateau (∼ 300 ms) gates modification of inputs active within a seconds-long eligibility window. **b**, Minimal model comprising a binary synaptic state *w*, an input pattern *x* and a shared scalar gate activation *f* (*w, x*). An accepted update replaces *w* with *x*, switching every mismatched synapse; otherwise, *w* remains unchanged (filled upward triangles, potentiation; open downward triangles, depression). **c**, The jump- or-stay rule defines a Markov chain on {±1}^*n*^ (three dimensions shown schematically). For the self-overlap gate illustrated, sufficiently similar inputs produce an accepted state-changing update, whereas dissimilar inputs leave the state unchanged. **d**, Anatomical interpretation of the plateau gate. The instructive gate integrates attenuated proximal drive *η* ⟨*w, x*⟩ with distal drive ⟨*v, x*⟩ ; the sigmoid maps their sum to gate activation, and an update is accepted above the plateau threshold *κ*. The self-overlap gate depends only on the proximal similarity ⟨*w, x*⟩.

To formalize these event-triggered properties, we represent the synaptic state of a neuron by a binary vector *w* ∈ {±1}^*n*^, each of whose *n* components denotes a weak or strong synapse. On trial *t*, an independently sampled input pattern *x*(*t*) ∈ {±1}^*n*^ drives the one-step plasticity update

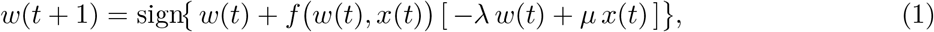

where sign acts coordinate-wise, 0 ≤ *λ, µ* ≤ 1, and 0 ≤ *f* ≤ 1 is the trial-specific scalar gate activation shared by all synapses. The parameters *λ* and *µ* quantify, respectively, the removal of the previous state and the imprinting of the current input. This minimal rule coordinates potentiation and depression within a common temporal window, with the direction determined by the current weight and input pattern, but idealizes every accepted update as a wholesale rewrite. The BTSP-inspired continual-recognition network below retains state-dependent credit assignment while replacing wholesale copying with a fuller timing-dependent potentiation and depression rule.

One-shot plasticity therefore defines a gated jump process on the set of plastic synaptic states (Fig. 1c). Because the inputs are independent across trials and equation (1) depends only on *w*(*t*) and *x*(*t*), the distribution of the next state depends on the past only through the current state. The common scalar gate gives each transition a jump-or-stay form: away from ties, the coordinate-wise update reduces to the binary alternative (Fig. 1b)

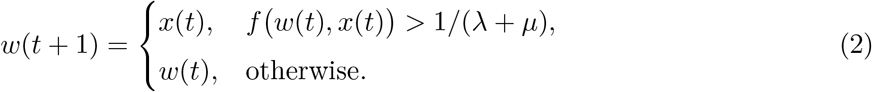

Each arriving input pattern therefore proposes a new plastic synaptic state. If the plastic event is accepted, all mismatched synapses switch and the accepted pattern is written as *w*(*t* + 1) = *x*(*t*); otherwise, the current state remains unchanged. We refer to this collective transition as the *jump- or-stay* reduction. It applies to any scalar gate and defines the transition structure analyzed below (proof in Supplementary Note 1). An accepted update applies the wholesale operation *w* ← *x*, which is the minimal learning rule. Graded alternatives such as partial rewrites or per-synapse failures, closely related to synaptic consolidation theory^39,40^, alter the distribution of resulting states while preserving the Markov description, provided that they introduce no additional history dependence.

Distinct biological organizations of the plastic and instructive pathways lead to distinct functional forms of the gate activation *f* . Each form determines event occurrence for each input and, together with the input statistics, the probabilities of acceptance and state change. We analyze two gate families (Fig. 1d). For the *self-overlap gate*, defined by the overlap between the plastic synaptic state and the current input (Hebbian-like self-addressing), only the plastic pathway contributes, and the gate activation increases with that state–input similarity, *f* = *σ*_g_(⟨*w, x*⟩ */σ*_0_ − *θ*_0_), where *σ*_g_ denotes the logistic function. For the *instructive gate*, an anatomically segregated distal synaptic weight vector *v* ∈ {±1}^*n*^ provides an additional contribution. Plateau initiation in CA1 pyramidal neurons reflects nonlinear integration of proximal Schaffer collateral input from CA3 and distal entorhinal input in the apical tuft^2,21^. We represent these contributions by the proximal plastic weights *w* and the distal instructive weights *v*, respectively:

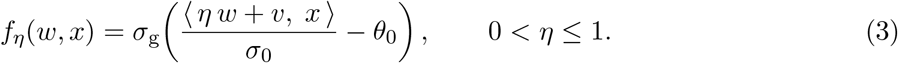

The integrated drive in equation (3) decomposes as ⟨*ηw* + *v, x*⟩ = *η*⟨*w, x*⟩ + ⟨*v, x*⟩ . Under this morphological interpretation, ⟨*w, x*⟩ is the integrated proximal input, attenuated by *η* as it propagates to the distal dendritic region in which the plateau is generated, whereas ⟨*v, x*⟩ is the locally integrated distal input. Thus, *η* phenomenologically represents the attenuation of the proximal contribution at the distal compartment, with *η* = 1 denoting the unattenuated limit. Here, *v* denotes the distal synaptic weight vector, which is held fixed over the timescale analyzed, while the corresponding instructive drive ⟨*v, x*(*t*) ⟩ varies across trials with the incoming pattern *x*(*t*), consistent with experience-dependent distal activity observed in vivo^22^.

In the active regime *λ* + *µ >* 1, the parameters *λ, µ, σ*_0_ and *θ*_0_ enter the event-acceptance criterion only through the *effective plateau threshold* 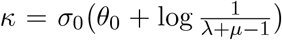 . Consequently, an update is accepted precisely when the total dendritic drive exceeds *κ* (Fig. 1d). The minimal model is therefore specified by the gate family, *κ*, the attenuation *η* for the instructive gate, and the statistics of the input stream.

### A single alignment variable governs memory and synaptic turnover

Applied to the BTSP-like instructive gate, the gated jump process reduces the synaptic dynamics to a single order parameter: pathway alignment. In the match-count convention, *m* = # {*i* : *w*_*i*_ = *v*_*i*_} counts the coordinates on which the plastic weights *w* match the instructive weights *v* (Fig. 2a), and the match fraction *m/n* measures this alignment within their shared input-feature basis. Thus *m/n* = 1 denotes perfect pathway agreement, whereas *m/n* = 1*/*2 is the chance-level baseline. For unstructured inputs instantiated by independent uniform signed patterns, permutation symmetry makes states with the same *m* statistically equivalent. The 2^*n*^ plastic synaptic states therefore project exactly onto *n* + 1 alignment levels (Fig. 2a), so this single variable characterizes both acquisition and long-term alignment dynamics (Supplementary Note 1).

**Figure 2:**
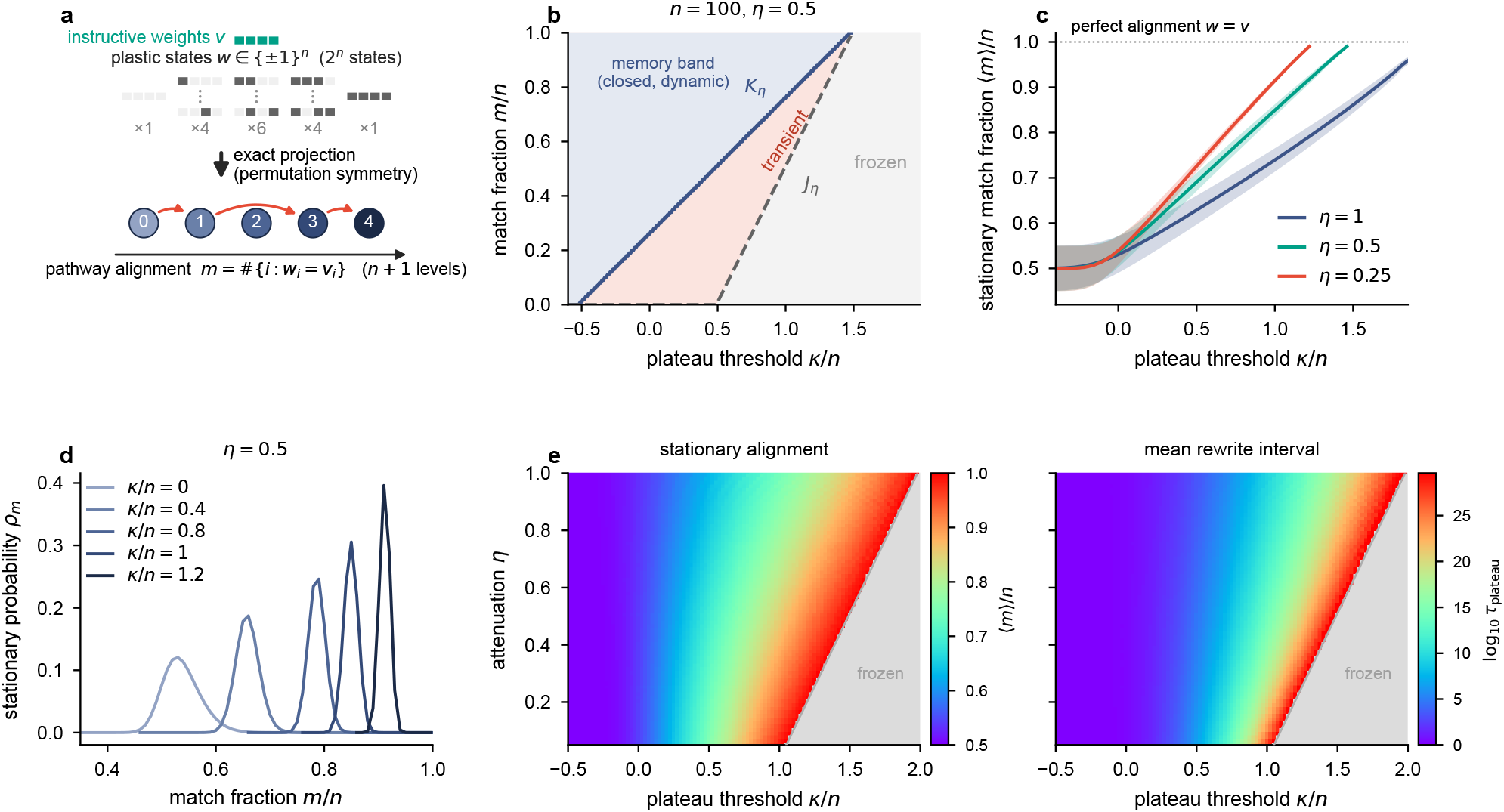
A single alignment variable governs memory and synaptic turnover. **a**, For independent uniform signed inputs, permutation symmetry projects the 2^*n*^ plastic synaptic states exactly onto *n* + 1 classes indexed by alignment *m* = # *i* : *w*_*i*_ = *v*_*i*_ (shown for *n* = 4); multiplicities are indicated below each class. **b**, Frozen and transient regions and the closed, dynamic memory band in the (*κ/n, m/n*) plane (*η* = 0.5). Accepted state-changing updates transfer transient states into the memory band; all states are frozen for *κ/n* ≥ (1 + *η*) − 2*η/n*. **c**, Exact stationary mean alignment (curves) and standard deviation (shading) as functions of *κ/n* for the indicated attenuation factors; the dotted line denotes perfect alignment. **d**, Exact stationary alignment distributions for *η* = 0.5 and the indicated thresholds. **e**, Exact stationary mean alignment (left) and mean rewrite interval log_10_ *τ*_plateau_ (right) over plateau threshold and attenuation, where *τ*_plateau_ is the stationary mean interval between accepted rewrite operations. Gray denotes the fully frozen parameter region. In **b**-**e**, *n* = 100

This reduction permits the complete finite-size dynamics and alignment statistics to be determined (Fig. 2b,c). The phase diagram contains three possible regimes. Poorly aligned states are *frozen*, because when the proximal plastic and distal instructive pathways are too different, their integrated drives can never produce an accepted update; intermediate states are *transient*, with each accepted state-changing update producing a one-shot jump into the memory band; and the *memory band* is closed but dynamic, with recurring accepted updates moving the weights among states within the band.

The reduced chain gives the transition and waiting statistics, together with the stationary mean, variance, and distribution of *m*, exactly. The plateau threshold *κ* controls both aspects: increasing it shifts the regime boundaries toward higher alignment, raises the stationary mean and reduces its dispersion, but eventually freezes plasticity. At the circuit level, inhibitory gating of dendritic input integration ^46,47^ motivates treating stronger inhibition as a higher effective plateau threshold *κ*. The theory therefore predicts an inhibition-dependent trade-off between plateau-recruitment frequency and fidelity of the instructed representation.

Stationarity does not imply a fixed plastic synaptic state (Fig. 2d). For *η* = 0.5, increasing *κ/n* from 0 to 1.2 shifts the exact stationary mean from ⟨*m*⟩ */n* = 0.537 to 0.912, yet every memory-band distribution shown has a mean below perfect alignment and is not concentrated at *m* = *n*. In this range, accepted state-changing updates continue to replace *w* while the distribution of its alignment with *v* remains stationary: microscopic synaptic turnover continues while coarse-grained macroscopic memory fidelity is preserved. Increasing *κ* reduces this turnover by shifting and narrowing the distribution; perfect alignment is approached only as state-changing accepted updates disappear near the freezing boundary. This coexistence of stable functional coding and continuing microscopic turnover provides a candidate mechanism for hippocampal representational drift^48,49^; recent trial-by-trial analyses further implicate rare, continuing BTSP events in this process ^50^.

The plateau threshold *κ* and attenuation factor *η* jointly control memory fidelity and kinetics (Fig. 2e). We define the *mean rewrite interval τ*_plateau_ as the stationary mean interval between accepted plastic events. Here, a rewrite denotes an accepted update operation and includes the idempotent no-op when *x* = *w*. Within the active region, increasing *κ* produces higher alignment and a much longer *τ*_plateau_; at positive thresholds, decreasing *η* has similar effects but induces freezing sooner. Microscopic state persistence is therefore kinetic: rare accepted updates slow synaptic turnover without requiring permanent stabilization of individual synapses. This mechanism complements our previous finding that BTSP repeatedly reconstitutes sustained CA1 place fields rather than permanently stabilizing the initially modified synapses^51^.

### The instructive pathway directs what can be learned from input structure

Natural input streams contain means, correlations and recurring patterns. Prior work showed that correlations reshape associative-memory storage and that novelty-facilitated plasticity can limit overassociation of correlated inputs^52,53^. The central question here is how this statistical structure interacts with the instructive pathway to determine the stationary plastic representation (Fig. 3a). The jump-or-stay reduction separates the problem into two levels. For any full-support input ensemble, the gate parameters (*v, η, κ*) fix the exact support of the dynamics—the frozen states, transient states and memory-band states—independently of the probabilities assigned to individual inputs. Those probabilities then determine the flow among the allowed states. Input structure thus controls how frequently different input patterns are proposed, whereas the instructive pathway biases which proposals produce accepted updates and are written into the plastic synaptic state. Over time, this acceptance bias shapes the stationary plastic representation (Supplementary Note 2).

**Figure 3:**
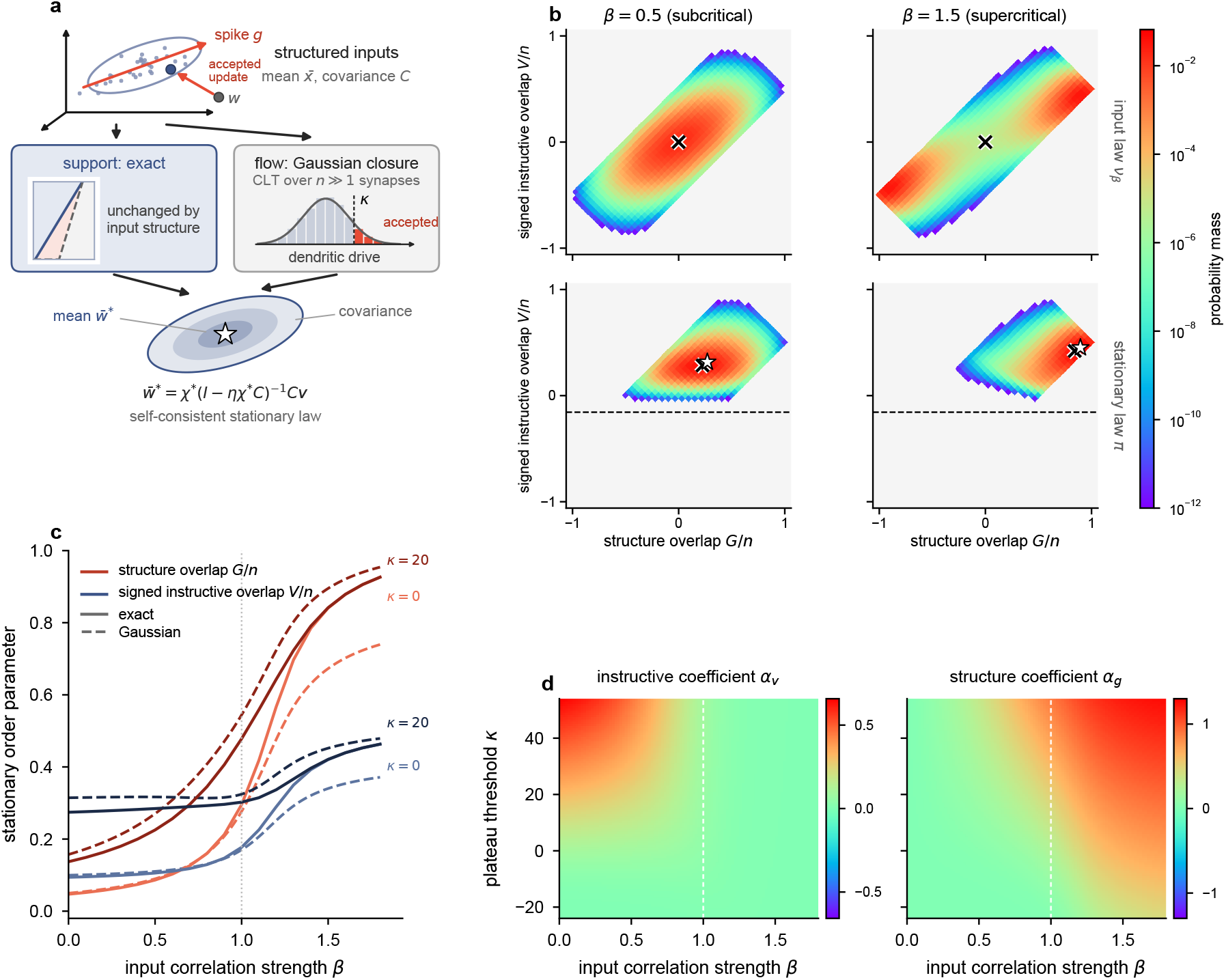
The instructive pathway directs what can be learned from input structure. **a**, Two-level reduction combining the exact support of the gated state-space dynamics with a Gaussian closure for the self-consistent stationary mean and a linear-noise approximation for the stationary covariance. **b**, Correlated bimodal Curie–Weiss input law (top) and exact stationary plastic-weight distribution under instructive gating (bottom), projected onto the structure overlap *G/n* and signed instructive overlap *V/n*, on opposite sides of the thermodynamic-limit critical value *β* = 1 (*β* = 0.5 and 1.5). In the bimodal regime, the instructive pathway selects the mode aligned with *v*. Crosses, exact means; stars, Gaussian predictions; dashed line, memory-band boundary; color, probability mass on a logarithmic scale. **c**, Exact (solid) and Gaussian (dashed) stationary means of *G/n* (red) and *V/n* (navy) as functions of *β* for *κ* = 0 and 20; dotted line, thermodynamic-limit critical value *β* = 1. **d**, Gaussian-resolvent coefficients in 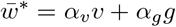 across the (*β, κ*) plane; dashed line, thermodynamic-limit critical value *β* = 1. In **b**–**d**, *n* = 64, *η* = 0.5 and *g*^⊤^*v/n* = 0.5; in **b**, *κ* = 20.

The uniform-input calculation above is the unstructured case, instantiated by independent uniform signed patterns. We isolated structured inputs using a correlated bimodal Curie–Weiss/Mattis ensemble ^54^, in which the correlation strength *β* controls the tendency of inputs to align with the structured directions ±*g* (Fig. 3b). Its covariance consists of an isotropic background plus a tunable rank-one component along ±*g*, providing a controlled analogue of spiked-covariance models in high-dimensional statistics^55,56^. The thermodynamic-limit critical value is *β* = 1; at the finite size considered here, the input law instead undergoes a smooth crossover from a single broad cloud to a distribution concentrated around the two modes ± *g*, while retaining zero mean. An exact projection onto *V* = *w*^⊤^*v* = 2*m* − *n*, whose normalized form *V/n* is the signed instructive overlap, and the structure overlap *G* = *w*^⊤^*g* reveals how the instructive gate resolves this symmetry. For *g*^⊤^*v/n* = 0.5, *η* = 0.5 and *κ* = 20, increasing *β* from 0.5 to 1.5 shifts the exact stationary mean from (*G/n, V/n*) = (0.225, 0.285) to (0.842, 0.421). At *β* = 1.5, on the supercritical side of the thermodynamic-limit critical value, the stationary law concentrates on the +*g* mode, which is more closely aligned with *v*, rather than averaging the two input modes. If the instructive vector is more closely aligned with the −*g* mode, the stationary law instead concentrates there (Supplementary Fig. S1). The instructive pathway therefore selects the component of input structure that is congruent with it, rather than simply overwriting that structure.

To approximate this interaction without enumerating the plastic synaptic state space, we Gaussianized the total dendritic drive *S*_*w*_ = (*ηw*+*v*)^⊤^*x* (Fig. 3c). This scalar combines the attenuated proximal contribution *ηw*^⊤^*x* with the distal instructive contribution *v*^⊤^*x*. Approximating its fluctuations as Gaussian while matching their exact first two moments converts event occurrence into a tractable tail probability. The resulting self-consistency relation approximates the stationary mean, and a subsequent linear-noise expansion provides an approximation to the stationary covariance. Against the exact Curie–Weiss chain, it captures the qualitative evolution of both *G/n* and *V/n* with *β*, their increase at the higher plateau threshold, and the growing influence of input structure across the finite-size crossover around *β* = 1. Quantitative discrepancies in the strongly correlated regime are analyzed in Supplementary Notes 2 and 3.

The stationary Gaussian closure gives the explicit resolvent

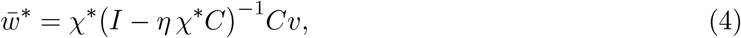

where *χ*^∗^ is fixed self-consistently by the plateau gate. For the Curie–Weiss covariance this reduces to 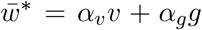, directly separating the instructive and input-derived components; Fig. 3d maps these formal coefficients. Within the admissible branch at the positive instructive–structure overlap shown, *α*_*v*_ is largest for weak input correlations and high *κ*, whereas *α*_*g*_ rises sharply as the input becomes bimodal and remains positive, selecting the mode which aligns better with *v*. The magnitude of this selection depends strongly on both *κ* and *g*^⊤^*v/n*: when the pathway overlap is zero, symmetry enforces *α*_*g*_ = 0 despite a bimodal law, whereas reversing the overlap reverses the selected structural component (Supplementary Fig. S1). Biologically, the distal instructive pathway therefore specifies which correlated proximal input ensemble is preferentially consolidated, while the plateau threshold controls the selectivity of that consolidation^21,22^.

### A BTSP-inspired network sustains continual recognition memory

We evaluated a BTSP-inspired continual-recognition network using the online protocol of Fig. 4a (Methods). Exact-*k* sparse binary patterns with coding fraction *k/d* = 0.05 arrived one per time step; after an initial novel segment, each subsequent pattern was either novel or, with probability 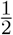, a repeat of the pattern presented *R* steps earlier, unless that pattern was itself a repeat (each pattern was thus repeated at most once). The network reported *novel* or *familiar* before plasticity acted on the current pattern. Its continuous familiarity score *y* then suppressed plasticity, a distinct mechanism that we call *familiarity-dependent plasticity suppression* (or recognition feedback; Methods). We defined the lag capacity *R*_0.90_, a retention-lag measure rather than a classical pattern-load capacity, as the largest tested repeat lag whose independently optimized network retained held-out protocol-weighted accuracy of at least 0.90.

**Figure 4:**
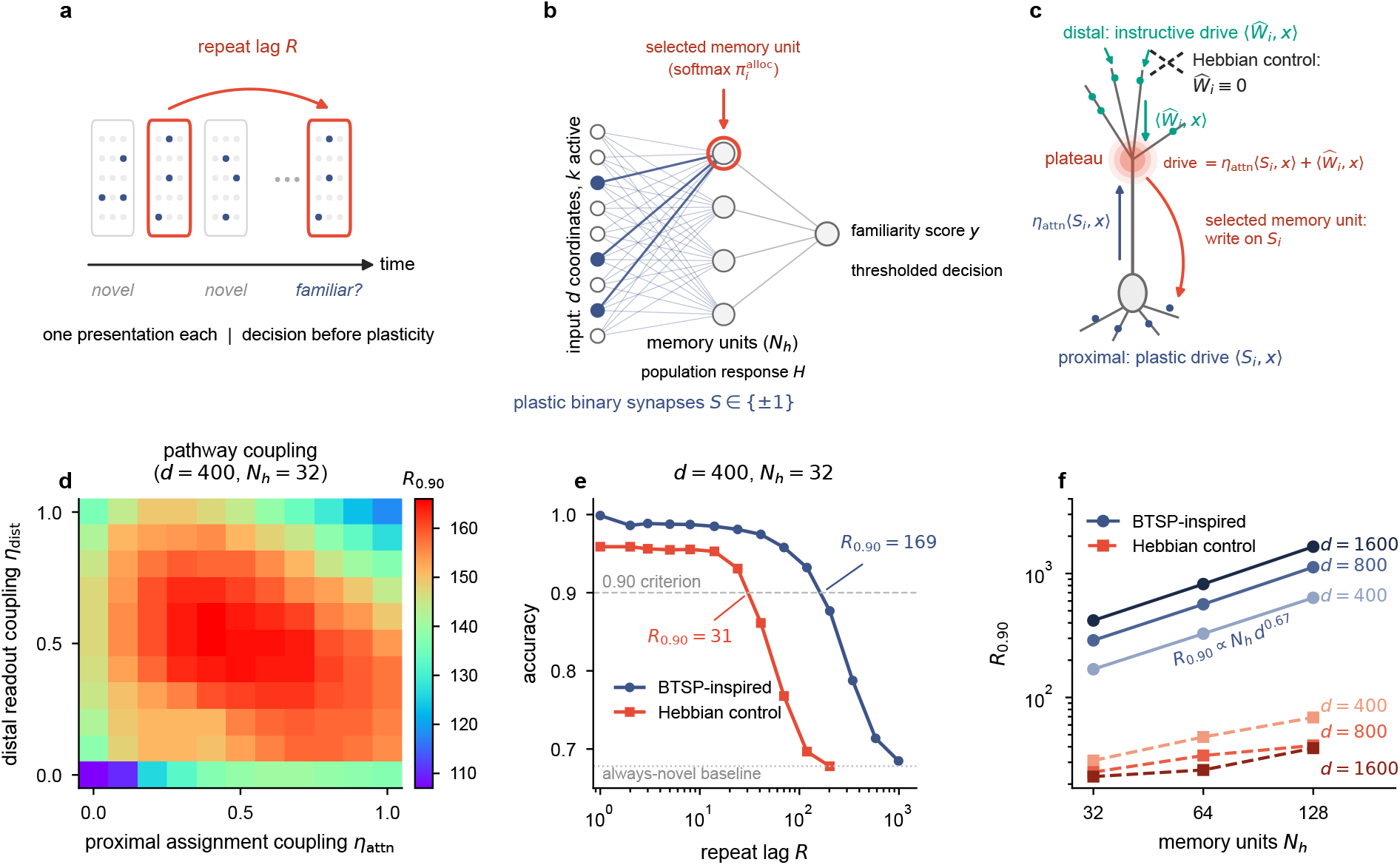
A BTSP-inspired network sustains continual recognition memory. **a**, Continual recognition task. A decision is made for each sparse binary pattern before plasticity; familiar trials repeat a pattern after repeat lag *R*. **b**, Continual-recognition network with a binary plastic memory matrix 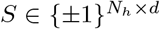, population response, and continuous familiarity score. Exact normalized softmax weights 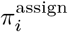 assign one-shot writes; the large optimized assignment gain makes one memory unit effectively the selected memory unit. **c**, The BTSP-inspired rule combines distal instructive drive 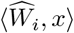 and proximal plastic drive *η*_attn_ ⟨*S*_*i*_, *x*⟩ to determine the selected memory unit. The Hebbian control removes the instructive pathway 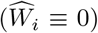 and uses independently balanced depression. **d**, Capacity *R*_0.90_ of the BTSP-inspired network versus proximal assignment coupling (*η*_attn_) and distal readout coupling (*η*_dist_). All other parameters are re-optimized for each condition. **e**, Per-lag optimized held-out protocol-weighted accuracy envelope for the BTSP-inspired network and Hebbian control. Their respective lag capacities are *R*_0.90_ = 169 and 31; dashed line, 0.90 criterion; dotted line, always-novel baseline. **f**, Lag capacity versus network width and input dimension. In **d**,**e**, *d* = 400 and *N*_*h*_ = 32.

The continual-recognition network has a binary plastic memory matrix 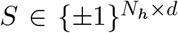 that connects the input to *N*_*h*_ memory units (Fig. 4b). Their memory-unit responses *h*_*i*_ form the population response *H*, from which the continuous familiarity score *y* and thresholded decision are obtained. Each memory unit *i* corresponds to a plastic row *S*_*i*_ and a fixed instructive row 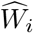, which play the respective roles of the single-neuron vectors *w* and *v*. The input dimension *d* replaces *n*, and the recognition task uses sparse binary patterns with exactly *k* active input coordinates, unlike the dense signed patterns used for the exact single-neuron reduction.

On every presentation, the plastic and instructive drives generate exact normalized softmax credit-assignment weights 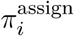. The full model retains this distributed softmax definition, but the assignment gain *g*_2_ was large enough in every optimized numerical network to concentrate the weights on the maximal-score memory unit. We therefore use *selected memory unit* throughout. The population model has no plateau threshold *κ*: *g*_2_ controls the concentration of credit assignment, while familiarity-dependent plasticity suppression separately scales the one-shot write.

The two learning rules differed in their credit assignment mechanisms and associated depression rules (Fig. 4c; Methods). In the BTSP-inspired network, drive 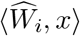 from fixed distal instructive weights was combined with proximal plastic drive *η*_attn_ ⟨*S*_*i*_, *x*⟩ to determine the selected memory unit, while the one-shot write modified the plastic matrix *S* but not the fixed instructive matrix 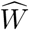. The Hebbian control removed the instructive pathway from the assignment rule and determined the selected memory unit through its current plastic drive^24,57,58^. The architectures were matched in plastic storage budget, binary plastic synapse type, one-shot-write budget, readout form, protocol, and optimization procedure; all free parameters were optimized independently for each condition.

The BTSP-inspired architecture permits reciprocal coupling between the distal instructive and proximal plastic pathways (Fig. 4d). The proximal assignment coupling factor *η*_attn_ is the network analogue of the single-neuron attenuation factor *η*, whereas the distal readout coupling factor *η*_dist_ sets the instructive contribution to the population response. Distal readout coupling alone raised *R*_0.90_ from 107 to 147, and proximal assignment coupling alone raised it to 140. Joint coupling produced a broad optimum, *R*_0.90_ = 166 at *η*_attn_ = 0.4 and *η*_dist_ = 0.5, that approximately recovered the capacity of the freely optimized network; pushing either contribution beyond its optimum instead degraded capacity. Intermediate optimal coupling implements partial, rather than complete, separation of credit assignment from plastic storage. With all free parameters optimized independently at each repeat lag, the BTSP-inspired network’s per-lag optimized accuracy envelope declined smoothly with repeat lag (Fig. 4e): at *d* = 400 and *N*_*h*_ = 32, held-out protocol-weighted accuracy decreased from 0.999 at *R* = 1 to 0.877 at *R* = 202, crossing 0.90 at *R*_0.90_ = 169 before approaching the always-novel baseline of 0.678 at longest tested repeat lags.

Input dimension revealed the clearest scaling difference between the BTSP-inspired network and Hebbian control: at every tested width, increasing *d* from 400 to 1600 raised the capacity of the BTSP-inspired network by about 2.5-fold but lowered the capacity of the Hebbian control (Fig. 4f). Across the tested grid *d* ∈ {400, 800, 1600} and *N*_*h*_ ∈ {32, 64, 128}, the capacity of the BTSP-inspired network grew with both dimensions, following the empirical finite-grid relation *R*_0.90_ ∝ *N*_*h*_*d*^0.67^ (*R*^2^ = 0.996), whereas the capacity of the Hebbian control increased only sublinearly with *N*_*h*_. At *d* = 400 and *N*_*h*_ = 32, *R*_0.90_ = 169 for the BTSP-inspired network versus 31 for the architecture-matched Hebbian control (Fig. 4e); across the grid, this advantage widened from 5.5-fold to 42-fold. We next use the reduced theory to examine the mechanisms underlying this divergent scaling.

### The reduced theory predicts memory in the full network model

We next applied the gated jump process to the full network model of Fig. 4. Mapping BTSP-inspired network architecture and learning rule onto the corresponding stochastic process yielded quantitative predictions of recognition performance (Fig. 5).

**Figure 5:**
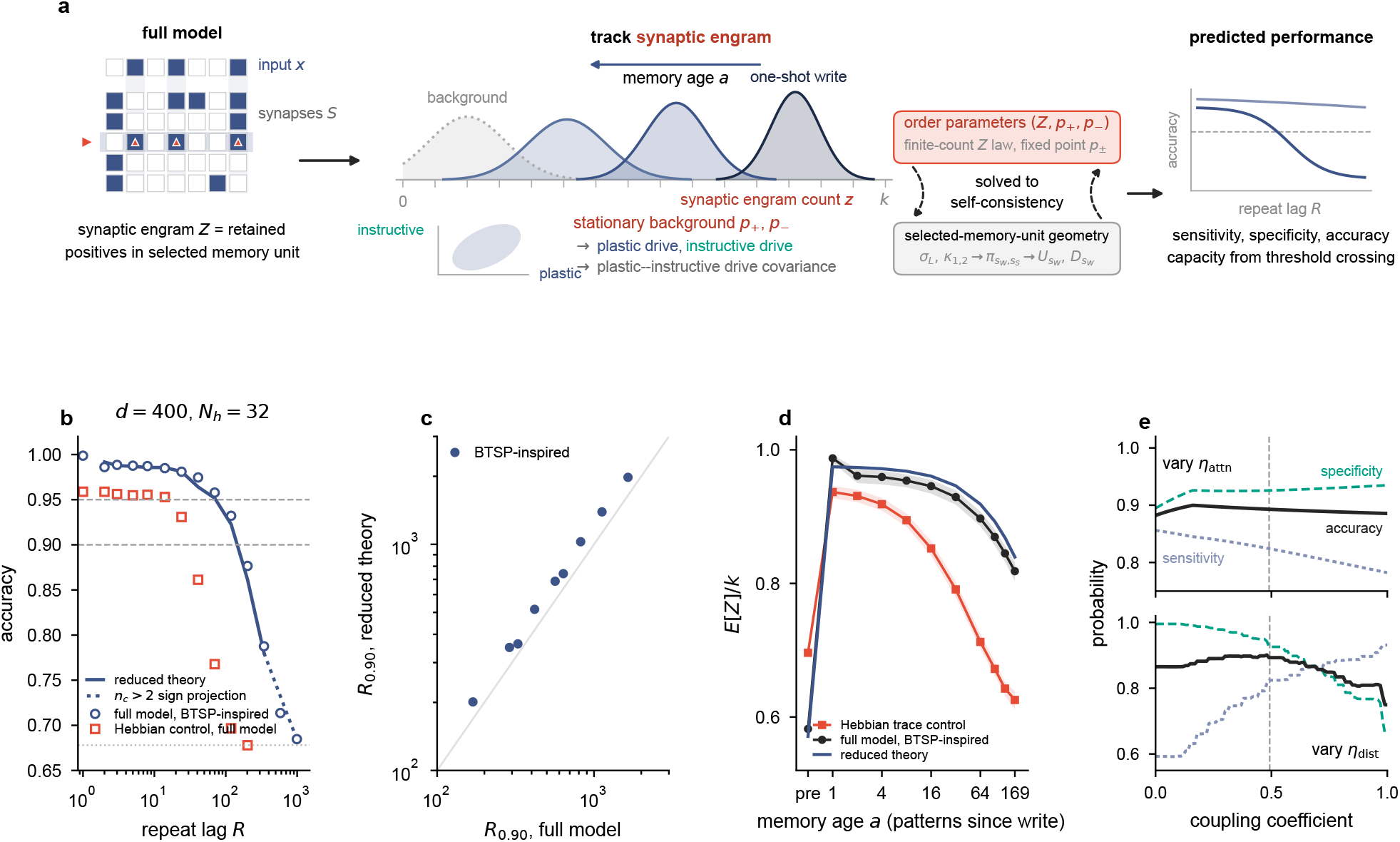
The reduced theory predicts memory in the full network model. **a**, The reduced theory tracks the synaptic engram *Z* and the stationary background (*p*_+_, *p*_−_), which fix the moments and covariance of the plastic and instructive drives. Order parameters and selected-memory-unit geometry (assignment-logit scale *σ*_*L*_, score moments *κ*_1,2_, enrichment 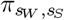, transition currents 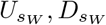; Supplementary Note 4.2) are solved self-consistently to predict response distributions, accuracy and capacity. **b**, Reduced theory for the BTSP-inspired network (solid, validated domain; dotted, sign projection for deep cascades with *n*_*c*_ *>* 2) and held-out full-model simulations of the BTSP-inspired network and Hebbian control (symbols) versus repeat lag. Dashed lines, protocol-weighted accuracy 0.90 and 0.95; gray dotted line, always-novel baseline. **c**, Lag capacity *R*_0.90_ of the BTSP-inspired network, optimized in the reduced theory versus that simulated in the full model for the nine network configurations in Fig. 4f; diagonal, identity. Geometric-mean prediction-to-simulation ratio, 1.20. **d**, Normalized memory-trace strength E[*Z*]*/k* from pre-write to memory age *a* = 169. Black, full-model simulation of the BTSP-inspired network; navy, reduced theory (root-mean-square error, 0.017); red, rate-and-depth-matched Hebbian trace control, with remaining parameters optimized at its own *R*_0.90_ = 31. Shading, pointwise 95% hierarchical-bootstrap intervals. **e**, Predicted novel-item specificity (teal, dashed), familiar-item sensitivity (slate, dotted), and protocol-weighted accuracy (black) at a fixed readout threshold while varying only proximal assignment coupling (top) or distal readout coupling (bottom); gray dashed lines, optimized coupling coefficients. In **b**,**d**,**e**, *d* = 400 and *N*_*h*_ = 32; in **d**, *a* = 169, and in **e**, *R* = 169.

The reduced theory extends the gated jump process from individual weights to the continual-recognition network in four stages (Fig. 5a; Methods; Supplementary Note 4). First, proximal plastic and distal instructive drives jointly determine the selected memory unit for each one-shot write. Second, potentiation and depression establish a stationary background summarized by *p*_+_ and *p*_−_, the probabilities that a plastic synapse is in the positive state conditional on a positive or negative instructive weight, respectively. The single-neuron quantities *m/n* and *V/n* express pathway alignment in match-fraction and signed-instructive-overlap conventions. In the full network, the corresponding statistic is the plastic–instructive correlation *c* = *p*_+_ − *p*_−_, whereas the separate plastic bias is *b*_*S*_ = *p*_+_ + *p*_−_ − 1. Conditional occupancies, Gaussian assignment-score statistics, and a categorical selected-memory-unit approximation close the stationary background self-consistently. Third, a tagged write creates the model-level synaptic engram *Z*, defined operationally as the number of active input coordinates retained as potentiated synapses in the selected memory unit. This synaptic-trace statistic is an analogue to the biological engram^59^. Later collisions at the same memory unit erode *Z*. We call E[*Z*]*/k* the normalized memory-trace strength, a predictor of the later familiar response, while the population response also depends on the other memory units (Fig. 5d). Finally, the tagged and background response distributions yield familiar-item sensitivity and novel-item specificity; weighting them by the continual-task priors instead gives protocol-weighted accuracy and lag capacity.

The reduced-theory prediction for the BTSP-inspired network’s per-lag optimized accuracy envelope closely tracked held-out simulations of the full model (Fig. 5b): across the 24 saved records at *d* = 400 and *N*_*h*_ = 32 within the validated theory domain, the mean absolute error was 0.010. Across the nine (*d, N*_*h*_) configurations, optimizing the reduced theory independently at each repeat lag preserved the simulated ordering and scaling of *R*_0.90_ but yielded an optimistic envelope: predicted capacities exceeded simulated values by a geometric-mean factor of about 1.20 (Fig. 5c).

To connect recognition performance to trace persistence, we tracked the normalized memory-trace strength E[*Z*]*/k* for prospectively tagged patterns, using the exact write-time softmax weights to average over the identity of the selected memory unit (Fig. 5d). At *d* = 400 and *N*_*h*_ = 32, the reduced theory reproduced the trajectory for the BTSP-inspired network through the pre-repeat state at memory age *a* = 169 with a root-mean-square error of 0.017. Between the immediate post-write state and memory age *a* = 169, this strength declined from 0.99 to 0.82 for the BTSP-inspired network, compared with a decline from 0.94 to 0.63 under a matched trace intervention of the Hebbian control. This Hebbian control matched the learning rate *η*_bin_ and cascade depth and used the same patterns and seeds; its remaining parameters were optimized at its own capacity, *R*_0.90_ = 31. The slower decay in the BTSP-inspired network is consistent with two complementary mechanisms for limiting interference: cross-time depression between successive plateau-defined activity patterns, related to a mechanism identified in CA3^20^, and credit assignment jointly determined by proximal and distal drive.

Furthermore, one-parameter reduced-theory sweeps clarify the interior coupling optimum of Fig. 4d (Fig. 5e; Supplementary Note 4.4, “How the two coupling coefficients act”). At *R* = 169, increasing proximal assignment coupling (*η*_attn_) allows current proximal weights to influence which memory unit gets selected. Specifically, memory units whose weights already match the input before the current write are more likely to generate a plateau and become the selected memory unit. This slows subsequent synaptic turnover and suppresses responses to novel inputs, raising novel-item specificity, but reduces potentiation headroom for writing a new pattern, weakening the initial familiar response and lowering familiar-item sensitivity. By contrast, distal readout coupling (*η*_dist_) leaves the synaptic engram and its decay unchanged while allowing stable instructive drive to bias the population somatic responses. Familiar inputs produce strong proximal and distal drive, increasing the somatic responses of the selected memory units and hence familiar-item sensitivity. However, distal readout coupling also broadens novel responses and lowers novel-item specificity; *η*_dist_ balances these effects to maximize recognition accuracy.

Finally, an assignment-opportunity concentration diagnostic revealed a further limitation of the Hebbian control (Supplementary Fig. S2). It measures expected, feedback-weighted update opportunities during continual presentation of patterns. The BTSP-inspired network maintained broad, nearly uniform relative assignment-opportunity shares across memory rows, whereas the Hebbian control progressively concentrated them onto a small subset. Concentration became more severe as *d* increased, so the declining effective assigned-memory-unit fraction may offset the potential storage benefit of higher-dimensional inputs. This provides a plausible, complementary explanation for why Hebbian capacity decreased rather than increased with *d* (Fig. 4f). Establishing the mechanism and its quantitative impact on capacity requires an extended reduced theory and remains future work. Switching the control to anti-Hebbian plasticity prevented this concentration by distributing one-shot writes broadly across memory rows and yielded capacity scaling with *d* and *N*_*h*_ comparable in form to that of the BTSP-inspired network, broadly consistent with earlier anti-Hebbian and optimized recognition-memory models ^13,15,60,61^ (Supplementary Fig. S3b,c). Nevertheless, its overall accuracy and capacity remained substantially lower than those of the BTSP-inspired network; at *d* = 400 and *N*_*h*_ = 32, it failed to reach 0.90 accuracy even at the shortest tested repeat lag (Supplementary Fig. S3a,b).

## Discussion

We developed a tractable theoretical framework connecting dendritic plateau event occurrence and synaptic modification to memory performance during continual learning. In the minimal model, the current plastic synaptic state and the instructive drive jointly determine whether a shared plateau event occurs, and that event in turn rewrites that plastic state (Fig. 1). For unstructured inputs instantiated by independent uniform signed patterns, permutation symmetry exactly groups the synaptic configurations by a single variable measuring pathway alignment between the plastic and instructive pathways, yielding the stationary plastic representation, synaptic turnover and the mean rewrite interval (Fig. 2). For structured inputs instantiated by correlated bimodal Curie–Weiss patterns, exact stationary analysis and a self-consistent Gaussian approximation to the summed dendritic drive show how the instructive pathway biases which component of input structure is incorporated into the stationary plastic representation (Fig. 3). At the network level, a reduced theory built on the same credit assignment logic predicts held-out protocol-weighted accuracy of the full model at saved parameter vectors, yields a per-lag optimized accuracy envelope and the resulting optimistic capacity, and reproduces the assignment-conditioned normalized memory-trace strength of a tagged synaptic engram through the pre-repeat state at memory age *a* = *R*. No closure coefficient was fitted to recognition accuracy (Fig. 5). Previous models showed that BTSP-like rules can support one-shot memory^43,44^, and familiarity networks with incremental plasticity have established capacity theories^13,15^. To our knowledge, however, a shared dendritic plasticity event whose occurrence depends jointly on the current plastic state and a distinct instructive pathway has not previously been analyzed together with its stationary transition and waiting-time statistics and quantitative continual-recognition performance.

The analysis identifies complementary mechanisms for memory persistence. First, plateau event occurrence stabilizes synaptic configurations kinetically. Because event occurrence depends jointly on the current plastic weights and the instructive drive, raising the plateau threshold makes accepted updates rare and lengthens the mean rewrite interval, while the instructive component biases the accepted updates toward the instructed representation. This mechanism is consistent with the reconstitution of sustained place fields by recurring BTSP events^51^. Second, the instructive pathway selects the component of structured input that is aligned with it (Fig. 3) and, in the network, provides an instructive drive for credit assignment that is not itself modified by each proximal one-shot write. Complete separation of credit assignment from storage was nonetheless not optimal. Distal readout coupling and proximal assignment coupling each produced a substantial capacity gain, and intermediate values of both coupling coefficients further increased the maximum observed capacity (Fig. 4d). The optimal architecture therefore separated assignment from storage only partially, combining functional specialization with controlled cross-pathway integration rather than relying on self-addressing credit assignment alone. A distinct feedback-mediated route to continual persistence uses retrievability-dependent stochastic rehearsal to consolidate memories^62^; here, recognition feedback instead suppresses new plasticity, and no deliberate replay is included.

A matched trace intervention asked whether the difference in persistence of memory-trace strength between the two rules reflects parameter differences alone. In the Hebbian trace control, the learning rate *η*_bin_ and cascade depth were set to the full-model values. Normalized memory-trace strength in the BTSP-inspired network still decayed more slowly than in the Hebbian control (Fig. 5d). The comparison does not isolate the remaining architectural and learning-rule differences, but the result is consistent with features of the BTSP-inspired rule that could limit interference: cross-time depression between successive plateau-defined activity patterns and selective credit assignment by the combined proximal and distal drive. Under the BTSP-inspired rule, only the currently and previously selected memory unit’s synaptic state is appreciably modified, whereas Hebbian depression of unselected memory units erodes their synaptic states on nearly every intervening step (Supplementary Note 4). A related cross-time mechanism supports memory in a recurrent CA3 model ^20^. In the capacity comparison, with each architecture at its own optimum, capacity in the BTSP-inspired network increased with both network width and input dimension, whereas Hebbian capacity showed weaker width dependence and declined at the largest input dimension (Fig. 4f).

Beyond these interference mechanisms, plastic-state-dependent credit assignment in the Hebbian control showed a further limitation. The assignment-opportunity concentration diagnostic showed that expected, feedback-weighted assignment opportunities progressively concentrated among a small subset of rows, reducing the effective allocated-memory-unit fraction as *d* increased, whereas the BTSP-inspired network maintained broad relative assignment-opportunity shares (Supplementary Fig. S2). Anti-Hebbian plasticity relieved this concentration and recovered capacity scaling with both *N*_*h*_ and *d* that was comparable in form to that of the BTSP-inspired network (Supplementary Fig. S3b,c), consistent with earlier anti-Hebbian models of recognition memory and related familiarity-capacity results ^13,15,60,61^. Its overall accuracy and capacity nevertheless remained substantially lower than in the BTSP-inspired network (Supplementary Fig. S3a,b). The BTSP-inspired network instead provides a different route to credit assignment: an additional stable instructive drive distributes one-shot writes broadly across selected memory units over time without anti-Hebbian plasticity. Previous works presenting similar separations of addressing from plastic storage use fixed distributed addresses ^63^ or a meta-learned addressing matrix^15^.

This two-pathway organization parallels the distinct entorhinal and CA3 contributions to plateau initiation and synaptic modification in CA1^21,22^. The theory linked the benefit of intermediate coupling between these pathways to distinct effects at the cellular and synaptic levels (Fig. 5e). When proximal plastic drive contributed to credit assignment, the selected memory unit was more likely to be one already responsive to the current pattern; its synapses turned over more slowly and novel patterns evoked weaker responses, but this also left less room to potentiate synapses during new encoding. Distal instructive drive instead contributed to the memory-unit response without changing the synaptic engram. It strengthened responses to familiar patterns but could also make some novel patterns appear familiar. Beyond these network-level effects, accepted updates continue to remodel plastic weights within the single-neuron model’s memory band, a closed but dynamic set of states, even though the pathway alignment distribution is stationary and remains below perfect alignment (Fig. 2c,d). Related theory has shown that stable collective memory can coexist with changing microscopic synapses^64^, although stability there resides in network dynamics rather than a stationary alignment distribution. The same gated jump process that maintains coarse-grained pathway alignment therefore produces synaptic turnover, a candidate mechanism for representational drift ^48–50^, although contributions from circuit reorganization remain outside the model.

The framework addresses a regime complementary to classical theories of synaptic memory. Palimpsest and cascade models ^33,36,37,39,40^ characterize the decay of memory signals under continual, distributed synaptic modification, and some allow one-shot or state-dependent transitions at individual synapses. Selective stochastic updating of bounded synapses has likewise been shown to extend memory lifetime^65^. Here, by contrast, eligibility is filtered through a shared dendritic event that the current weights help to trigger, so updates can be comparatively sparse and an accepted update can modify many eligible synapses in concert. The analysis therefore emphasizes transition kinetics and the geometry of reachable states rather than signal-to-noise propagation alone. BTSP motivates this regime, but the formalism is not restricted to a particular biological plasticity rule. Any process whose rules for event occurrence and updating depend only on the current synaptic state and input can be formulated as a gated jump process and represented by a Markov chain. Partial rewrites or synapse-specific failures change the transition kernel without altering this structure. The exact order-parameter reduction additionally requires appropriate symmetries, whereas the Gaussian closure requires a summed drive for which the Gaussian approximation is accurate.

The framework yields several experimentally testable predictions. First, plateau event occurrence should depend steeply on the combined distal and attenuated proximal drive, so that the probability of a plateau in response to a given input depends on the current synaptic state and not on the input alone. Second, if stronger inhibitory control of plateau recruitment raises the effective plateau threshold, the theory predicts higher pathway alignment and fewer accepted updates within the active regime, with freezing at sufficiently high inhibition. Third, for structured inputs, the distal instructive pathway should preferentially consolidate the correlated proximal ensemble most closely aligned with it. Finally, acquisition should be step-like at the single-cell level. In the minimal model, a recruitable neuron’s plastic synaptic state enters the memory band through a single plastic event rather than through gradual accumulation.

The present analysis assumes binary synaptic states, wholesale rewriting by accepted updates and an instructive weight vector that is fixed over the analyzed timescale, idealizing an instructive pathway that changes more slowly than the plastic weights, although its input-dependent activity varies across trials. The exact single-neuron results further assume temporally independent input draws, and the reduced theory treats prescribed repeats rather than general temporal correlations. Quantitative validation of the reduced theory is restricted to shallow metaplastic cascades and to the regime in which its Gaussian closures hold. Incorporating graded synaptic changes, synaptic decay, correlated experience streams, deeper metaplasticity and recurrent circuit dynamics constitutes a natural next step, and the separation between exact support geometry and closed probability flow specifies where each extension enters the theory.

Many approaches to continual learning in artificial systems combine gradual gradient updates with regularization, replay or architectural mechanisms that limit interference ^9,10,66^. The BTSP-inspired strategy examined here instead uses sparse one-shot writes, coordinated modification of eligible synapses and a partly distinct instructive pathway to help determine the selected memory unit for each write. Across the tested networks, this architecture produced a capacity advantage over the Hebbian control that increased from 5.5-fold to 42-fold while remaining amenable to quantitative analysis. These results show that one-shot plasticity is compatible with both persistent continual memory and analytical tractability. Plasticity in which the synaptic state helps select the events that modify it, combined with partial separation of credit assignment from memory storage, therefore offers a biologically motivated design principle for continual learning.

## Methods

### The gated jump process

The synaptic state is *w* ∈ {±1}^*n*^ and inputs *x*(*t*) arrive independently across time. Weights evolve by equation (1) with 0 ≤ *λ, µ* ≤ 1 and a scalar gate activation 0 ≤ *f* ≤ 1; sign acts coordinate-wise.

Because *f* is shared by all coordinates, either every mismatched coordinate flips or none does, giving the jump-or-stay reduction, equation (2); if *λ*+*µ* ≤ 1 the chain is frozen for any gate (Supplementary Note 1). In the active regime *λ* + *µ >* 1, for logistic gates *f* = *σ*_g_(*D/σ*_0_ − *θ*_0_) with drive *D*, the event-occurrence condition *f >* 1*/*(*λ* + *µ*) is equivalent to *D > κ* with 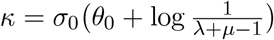; all results depend on (*λ, µ, σ*_0_, *θ*_0_) only through *κ*. The self-overlap gate uses *D* = ⟨*w, x*⟩; the instructive gate uses *D* = ⟨*ηw* + *v, x*⟩ with instructive weight vector *v* ∈ {±1}^*n*^ and attenuation 0 *< η* ≤ 1 (equation (3)); *η* = 1 recovers the unattenuated instructive gate as a special case.

### Exact order-parameter reduction (Fig. 2)

For independent uniform signed inputs and the instructive gate, the match count *m* = #{*i* : *w*_*i*_ = *v*_*i*_} defines an exactly lumped Markov chain. Conditional on *m*, write 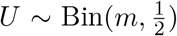 and 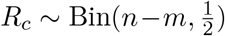 for the numbers of instructive-aligned coordinates confirmed and corrected by the input; the proposed next level is *ℓ* = *U* + *R*_*c*_ and the drive is *G*_*η*_ = 2(1 − *η*)*ℓ* + 4*ηU* (1 − *η*)*n* − 2*ηm*. The accepted-update matrix is

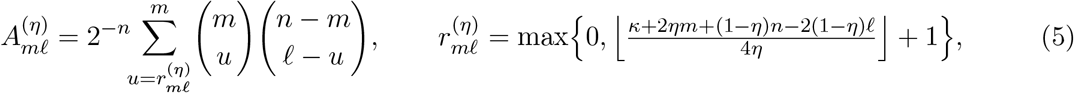

with event-occurrence probability 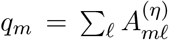 and held chain *Q* = *I* − diag(*q*) + *A*^(*η*)^. The zone edges are *K*_*η*_(*κ*) = max 0, ⌊[*κ* + (1 − *η*)*n* + 2*η*]*/*2⌋ + 1 (memory-band floor) and *J*_*η*_(*κ*) = max 0, ⌊[*κ* − (1 − *η*)*n*]*/*(2*η*)⌋ + 1 (edge of the frozen zone); levels *J*_*η*_ ≤ *m < K*_*η*_ have a geometrically distributed waiting time followed by one accepted state-changing update into the band. The memory band is closed but dynamic. On this band the stationary alignment law factorizes into jump statistics and dwell times: if *ξ* is the stationary law of the embedded accepted-update chain 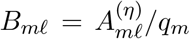, then *ρ*_*m*_ ∝ *ξ*_*m*_*/q*_*m*_. The full-state stationary law is uniform within each level. The stationary accepted-event rate is 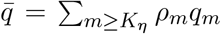, and Fig. 2e reports the mean rewrite interval 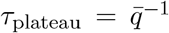. Here a rewrite is an accepted update operation *w* ← *x*, including an idempotent operation when *x* = *w*. Statements and proofs are given in Supplementary Note 1. For Fig. 2 the chain was solved exactly at *n* = 100 (log-domain binomial sums; power iteration to 10^−14^). For the self-overlap gate the alignment autocorrelation obeys E[*w*(*t*) | *w*(0)] = *α*^*t*^*w*(0) with 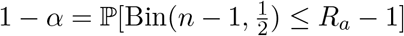, where *R*_*a*_ is the accepted Hamming radius.

### Structured inputs and the Gaussian closure (Fig. 3)

Let structured inputs be drawn from an ensemble *ν* on {±1}^*n*^ with mean 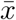 and covariance *C*. The jump graph *w* → *y* ⇐⇒ (*ηw*+*v*)^⊤^*y > κ* depends only on the support of *ν*; for full-support ensembles the frozen/transient/band geometry is identical to the uniform case (Supplementary Note 2). The stationary law on the band solves a directed balance equation and generically carries currents (no detailed balance). The Gaussian closure treats the drive *S*_*w*_ = (*ηw* +*v*)^⊤^*x* as Gaussian with the exact moments 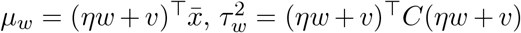, giving 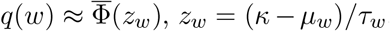, and the accepted-update mean 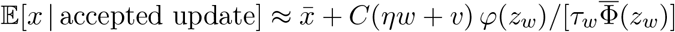. For balanced ensembles 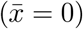, the representative-state closure of Supplementary Note 2 and zero approximate mean drift yield the self-consistent mean of equation (4), with 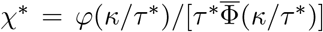 and 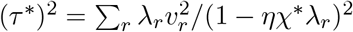 summed over the eigenmodes *λ*_*r*_ of *C*; the admissible branch requires *ηχ*^∗^*λ*_max_(*C*) *<* 1, binary bounds 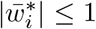, and the exact band floor 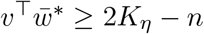. Under the frozen-coefficient and noise simplifications detailed in Supplementary Note 2, linearization about 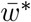 gives a discrete Lyapunov equation whose modal solution is a linear-noise approximation to the stationary covariance.

### The Curie–Weiss input ensemble

The solvable correlated family is 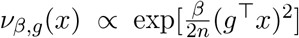 for a fixed pattern *g* ∈ {±1}^*n*^: a balanced, full-support ensemble whose inputs share a tunable similarity to ±*g*^54^ . Its covariance is *C* = (1 − *r*)*I* + *r gg*^⊤^ with the exact pair correlation *r*_*β,n*_ = (E_*β*_[(*g*^⊤^*x*)^2^] − *n*)*/*[*n*(*n* − 1)]. In the thermodynamic limit, the collective (Mattis) eigenvalue *λ*_*g*_ = 1 + (*n* − 1)*r*_*β,n*_ becomes *O*(*n*) for *β >* 1, with critical point *β* = 1; at finite *n*, including *n* = 64 here, this is a smooth crossover toward the two directions ±*g* (Fig. 3b). Exact stationary laws (Fig. 3b,c) were obtained with no sampling by a second exact projection: the signed permutations fixing *g* and *v* lump the chain onto the pair (*a*_+_, *a*_−_) counting matches with *g* inside the coordinate classes *g*_*i*_ = *v*_*i*_ and *g*_*i*_ = −*v*_*i*_ (equivalently onto (*G, V* ) = (*w*^⊤^*g, w*^⊤^*v*)), giving a finite orbit chain whose transition kernel is a hypergeometric tail sum weighted by the ensemble (Supplementary Note 3); its normalized left eigenvector with eigenvalue 1 is the exact stationary law. Panels used *n* = 64, *h* = *g*^⊤^*v* = 32, *η* = 0.5; the Gaussian predictions come from the scalar self-consistency above specialized to the two-eigenvalue covariance (Fig. 3c; dashed curves are drawn only where the branch is admissible).

### Continual recognition memory protocol

At each time step *t* ∈ {0, …, *T* − 1} a pattern *x*_*t*_ ∈ {0, 1}^*d*^ with exactly *k* active input coordinates (*k* = ⌊0.05 *d*⌋) is presented; the first *R* steps are forced novel, and each later step repeats the pattern from lag *R* with probability 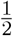 unless that pattern was itself a repeat, so that every pattern is repeated at most once (single repetition; repeats do not chain). The horizon is *T* = max(200, 20*R*), which pins the always-novel baseline accuracy at 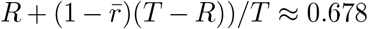 for *R* ≥ 10, where 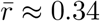 is the repeat fraction averaged over the *T* − *R* repeat-eligible steps (the per-step repeat probability relaxes to 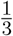 because a repeated pattern is never repeated again). The network makes its familiar/novel decision before its own plasticity for that step, and its continuous familiarity score controls plasticity through the familiarity-dependent plasticity-suppression multiplier described below; protocol-weighted accuracy is the fraction of correct reports over all steps and parallel streams. Reported accuracies are *held out*: each tuned record is evaluated on a separate input tensor generated with data seed 10,000 + *R*, never used during tuning, and averaged over 16 model state/event seeds, 2000, …, 2015. Lag capacity *R*_*θ*_ is the largest tested lag with held-out protocol-weighted accuracy ≥ *θ*, measured on a geometric lag ladder (*R* → max(round(1.7*R*), *R* + 1) from *R* = 1); each rung is a complete tune-then-test experiment, and bisection localizes the crossing between the last passing and first failing rung to adjacent integers. Because each lag is tuned independently, reported capacities are search-dependent lower envelopes: adjacent-integer bisection localizes the tested crossing but does not exclude downward bias from incomplete optimization.

### The BTSP-inspired continual-recognition network

The network holds a plastic sign matrix 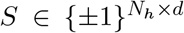 (optionally with metaplastic cascade depth *n*_*c*_^39^) and a fixed random instructive matrix 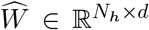 obtained by subtracting each row mean from an i.i.d. Rademacher matrix, without row normalization (fixed instructive seed 12,345). The cascade depths at the tuned optima are shallow at most rungs, although two of the three capacity-crossing optima at *d* = 1600 select *n*_*c*_ = 3. Records with *n*_*c*_ *>* 2 lie outside the formal theory domain; the two deep-cascade records in Fig. 5b are shown only as dotted, unscored sign projections. On input *x* with plastic drive *Sx* and instructive drive 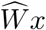, the memory-unit responses are 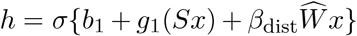; the normalized credit-assignment weights are 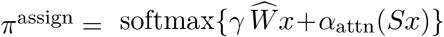 ; and the continuous familiarity score is *y* = *σ*{*b*_2_+*w*_2_ ∑_*i*_ *h*_*i*_}, thresholded at 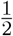. The slopes are 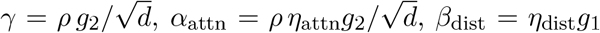 (the centered instructive rows have approximately 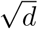 norm, matching the scale of the rows of *S*), with 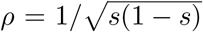 normalizing the instructive drive at coding fraction *s* = *k/d*, so *η*_attn_ and *η*_dist_ are dimensionless coupling coefficients: *η*_attn_ is the proximal assignment coupling and the network analogue of the framework’s attenuation *η*, whereas *η*_dist_ is the distal readout coupling. At every optimized operating point the assignment softmax is effectively in the hard-winner regime; we therefore call each memory unit with a maximal assignment weight a selected memory unit, while retaining the exact fractional weights in the full model.

Plasticity is event-driven and one-shot: per step, three plasticity passes are applied to assignment-weighted synapses—depression of the current pattern’s active input coordinates weighted by the *previous* assignment 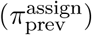, depression of the previous pattern’s active input coordinates weighted by the current assignment (*π*^assign^), then potentiation of the current pattern’s active input coordinates weighted by *π*^assign^—with probabilities *η*_dep_, *η*_dep_, *η*_bin_ and the structural tie *η*_dep_ = *η*_bin_*/*2. For *n*_*c*_ = 1, a delivered opposing event flips the readable sign and a matching event leaves it unchanged. For *n*_*c*_ *>* 1, opposing events flip and reset with a depth-dependent probability, whereas matching events can deepen the synapse without changing its readable sign. All network simulations fixed (`cascade_x`, *f*_pot_, *f*_dep_) = (1*/*2, 1*/*2, 1*/*2); under this symmetry, *n*_*c*_ = 2 has the same readable-sign and readout dynamics as *n*_*c*_ = 1. Every plasticity pass is additionally multiplied by the recognition-feedback multiplier *f*_fb_ = 1 − *y* computed from the current familiarity score. This familiarity-dependent plasticity suppression becomes stronger as the pattern appears more familiar, while credit assignment remains active. Rather than being drawn without bias, the readable plastic signs are initialized from the conditional stationary product-law closure of single-level binary, all-novel dynamics, conditioned on the instructive sign, 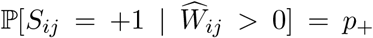 and 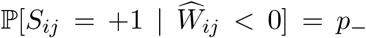. Here (*p*_+_, *p*_−_) is the fixed point of a self-consistent single-presentation closure of the potentiation and depression passes, solved for each parameter vector from (*η*_bin_, *η*_dep_, *η*_attn_, *g*_2_) together with the network shape and coding fraction (*p*_+_ = 0.54, *p*_−_ = 0.33 at the *R*_0.90_ optimum of Fig. 4e). For *n*_*c*_ *>* 1, these probabilities specify only the readable-sign marginal; cascade depths are initialized uniformly. Model-side instructive matrix, initialization and plasticity-event draws use counter-based Philox randomness, whereas input tensors are generated deterministically with the retained legacy NumPy stream. For fixed inputs and seeds, the resulting discrete state trajectories are bit-exact across software backends.

### The Hebbian control

The Hebbian control removes the instructive pathway: its credit assignment reads the plastic matrix itself, 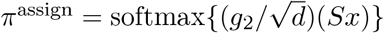, and *h* = *σ*{*b*_1_ +*g*_1_(*Sx*)}. Its three same-step plasticity passes are applied sequentially: LTD on inactive input coordinates with assignment weight 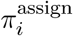 and probability *η*_bi_*/*[2(*Nh*− 1)]; LTD on active input coordinates with weight 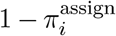 and probability *η*_bin_*η*_bin_/[2(*N*_*h*_ − 1)]; then LTP on active input coordinates with weight 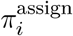 and probability *η*_bin_.

Following the general balance principle for associative matrix memories ^67^, the two LTD probabilities are derived here, not tuned, so that the expected depression event mass exactly balances the expected potentiation event mass. Integer-valued drives make ties among highest-assignment memory units frequent; tied selected memory units share the write. Everything else (readout, protocol and the recognition-feedback multiplier *f*_fb_ = 1 − *y*) is identical to the BTSP-inspired network. The plastic signs are likewise initialized with a bias from the stationary product-law closure of single-level binary, all-novel Hebbian plasticity passes. This closure has no instructive pathway to condition on and, at the balanced depression probabilities, depends only on (*k, N*_*h*_): ℙ [*S*_*ij*_ = +1] = *p* with *p* = 0.39 at (*d, N*_*h*_) = (400, 32) (0.36–0.44 across the grid). For *n*_*c*_ *>* 1, this probability likewise specifies only the readable-sign marginal; cascade depths are initialized uniformly.

### The anti-Hebbian control

For the anti-Hebbian control, the polarity of every Hebbian plasticity pass was reversed, reported in the anti-Hebbian convention as *η*_bin_ *<* 0: the 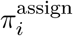 -weighted active-coordinate pass depressed its targets, whereas the 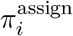 -weighted inactive-coordinate and 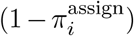 -weighted active-coordinate passes potentiated theirs at the same balanced probability magnitudes. Thus |*η*_bin_| is the underlying learning rate. All row and column masks, familiarity-dependent plasticity suppression, stationary sign initialization, cascade dynamics and the single-repeat protocol were retained. Because a familiar pattern’s stored synaptic modification lowers the memory-unit response, the production search used *w*_2_ *<* 0 and expressed the memory-unit and readout biases in scale-independent threshold coordinates: *b*_1_ = −*g*_1_*kθ*_1_, *w*_2_ = −*w*_2,mag_ and *b*_2_ = *w*_2,mag_(*N*_*h*_ − *m*_out_). Two analytic candidates seeded only the first rung of each arm; subsequent rungs used the usual neighboring-rung warm start. Because the *d* = 400 network did not reach 0.90 protocol-weighted accuracy, *R*_0.80_ was added for Supplementary Fig. S3. Apart from these stated exceptions, the anti-Hebbian campaign used the same ten-arm layout and adaptive TPE budget described below.

### Per-condition tuning

The BTSP-inspired network and Hebbian control were each run on ten *arms*: the nine grid configurations (*d, N*_*h*_) ∈ {400, 800, 1600} × {32, 64, 128} plus a dense lag sweep run to the always-novel baseline at (*d, N*_*h*_) = (400, 32) (the arm shown in Figs. 4e and 5b). For every arm and every ladder lag, all free parameters (*η*_bin_, *g*_1_, *g*_2_, *b*_1_, *b*_2_, *w*_2_, cascade depth, plus *η*_attn_, *η*_dist_ for the BTSP-inspired network) were tuned by Bayesian optimization (TPE^68^; adaptive budget up to 2,000 trials per rung, warm-started from the neighboring rung) on search data, then evaluated on held-out data as above. Both architectures therefore compete at their per-condition optima. Individual seeded evaluations are deterministic, but parallel TPE proposal sequences can depend on trial completion order; the saved per-rung records are authoritative. For Fig. 4d, (*η*_attn_, *η*_dist_) was fixed on an 11 × 11 grid over [0, 1]^2^, and the remaining seven parameters were re-optimized in every grid point and at every queried lag using the same protocol.

Finite-grid scaling exponents were obtained by ordinary least-squares regression of log *R* on log *N*_*h*_ and log *d*, with the coefficient of determination evaluated in log space. The fit for the BTSP-inspired network in Fig. 4f used all nine configurations; its fitted *N*_*h*_ exponent is 0.98. The anti-Hebbian *R*_0.80_ fit in Supplementary Fig. S3b used only the six configurations with *d* = 800 or 1600, as specified in the caption.

### Assignment-opportunity concentration metrics

To measure persistent differences in assignment opportunity among labeled memory units, a fixed network snapshot at learning time *t* was probed with *N*_*q*_ independent, read-only novel patterns. Let 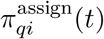 be the credit-assignment weight of memory unit *i* on probe *q*, and let *f*_*q*_(*t*) = 1 − *y*_*q*_(*t*) be the familiarity-dependent plasticity-suppression multiplier. The feedback-weighted assignment opportunity and relative assignment-opportunity share of memory unit *i* are

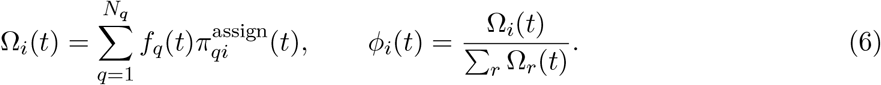

Without familiarity-dependent plasticity suppression, *f*_*q*_ = 1. We summarized the assignment-opportunity-share distribution using

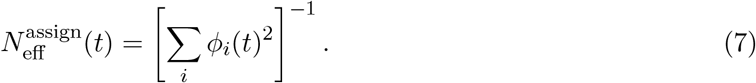

Here 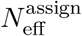 is the effective number of memory units available for credit assignment.

For Supplementary Fig. S2a,b, states from a representative *d* = 400, *N*_*h*_ = 32, *R* = 169 stream were retained as fixed network snapshots at checkpoints through presentation 8,192 and evaluated on a common bank of *N*_*q*_ = 2,048 independent exact-*k* novel probes. For panel c, each size configuration was evaluated at presentation 8,192 using *N*_*q*_ = 8,192 probes and 128 trajectories. The BTSP-inspired network used each configuration’s saved *R*_0.90_ record; the Hebbian control used gain and readout parameters from its own *R*_0.90_ record, with *η*_bin_ and cascade depth replaced by the corresponding values from the BTSP-inspired network, and both networks received the BTSP-inspired network’s *R*_0.90_ stream. Panel-c 95% intervals used 2,000 crossed-factor bootstrap replicates over data seed, state seed and stream. Supplementary Fig. S3c used the same representative stream and probe bank with the anti-Hebbian control’s own *R*_0.80_ record.

### Theory evaluation

The reduced theory (Supplementary Note 4) uses single-level binary synapses, exactly balanced binary instructive rows, a centered plastic credit-assignment drive, and a categorical selection that replaces the full model’s distributed softmax assignment weights with one selected memory unit. It also uses independent depression sets, disables familiarity-dependent plasticity suppression, and includes no deliberate replay of the tagged pattern. Independent Gaussian assignment scores are summarized by their scale *σ*_*L*_ and selected-score moments (*κ*_1_, *κ*_2_); a second-order Hermite projection gives the sign-conditioned coordinate enrichments and transition currents that close the stationary background (*p*_+_, *p*_−_). This pair also fixes the moments of the plastic and instructive memory-unit drives *a*_*i*_ and *v*_*i*_, including *C*_*av*_ = Cov(*a*_*i*_, *v*_*i*_). A tagged write creates the synaptic engram *Z*, the raw positive count among its *k* active input coordinates, resolved by instructive-sign class. Later interference is propagated with the finite collision count

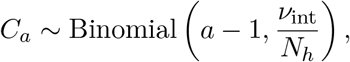

where *ν*_int_*/N*_*h*_ is the per-presentation collision rate. Finite generic- and tagged-memory-unit drive lattices are passed through the memory-unit sigmoid and combined with a shared nonnegative, two-moment gamma remainder to obtain novel-item specificity, familiar-item sensitivity, and balanced accuracy. No closure coefficient is fitted to balanced accuracy.

For comparison with the full model’s continual single-repeat campaign, the reduced theory fixes *ν*_int_ = 2*/*3 and treats familiar presentations as exact no-ops. This thins only the post-write collision rate, leaving the stationary background and immediate write unchanged. To match the full model’s protocol-weighted accuracy objective, theory specificity and sensitivity are weighted by the novel

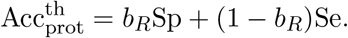

The saved-record comparisons use the empirical *b*_*R*_; reduced-theory optimization uses the exact finite-*T* protocol prior. The fixed thinning is a protocol approximation, not a fitted correction.

The record-level validation underlying Fig. 5b evaluates the theory at the parameter vector for each saved BTSP-inspired network. Of 27 saved *d* = 400, *N*_*h*_ = 32 records, *R* = 1 is omitted as a boundary case, 24 records with *n*_*c*_ ≤ 2 are formally scored, and the *R* = 583 and 991 records with *n*_*c*_ = 3 and 4 are shown only as dotted, unscored sign projections. The mean absolute error in protocol-weighted accuracy across supported records is 0.00965 (reported as 0.010).

Figure 5c instead uses an independent reduced-theory optimization at every queried lag, with *n*_*c*_ = 1. The search uses the full model’s wide parameter domains for (*η*_bin_, *g*_1_, *g*_2_, *b*_1_, *η*_attn_, *η*_dist_), profiles the invariant readout threshold Θ = − *b*_2_*/w*_2_ exactly, screens scrambled Sobol candidates with local refinements, and reranks finalists with the full finite-lattice/gamma calculation. The full model’s geometric ladder and adjacent-integer bisection are retained. Primary searches use 4,096 candidates for *d* ≤ 800 and 8,192 for *d* = 1600, four local refinements and 32 full-calculation finalists. Provisional failures are retried independently; final boundaries use at least 8,192 candidates and 64 full-calculation finalists. Across the nine configurations of the BTSP-inspired network, the geometric-mean reduced-theory/full-model *R*_0.90_ ratio is 1.201 and the mean absolute relative error is 20.18%. This is an optimistic independently optimized envelope, not the retention trajectory of one fixed classifier.

For a prospectively tagged pattern *p* at memory age *a*, with active set *A*_*p*_ = {*j* : *x*_*p,j*_ = 1}, the full model’s normalized memory-trace strength in Fig. 5d is the credit-assignment-weighted positive fraction

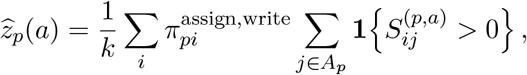

using each model’s assignment weights fixed at write time. The “pre” point precedes the tagged write, and *R*^−^ = 169 is the state immediately before lag-*R* classification and updating. This is the normalized memory-trace strength, not the causal engram contrast against the matched no-write control in Supplementary Note 4. The theoretical line is the corresponding E[*Z*]*/k* at the saved anchor with *ν*_int_ = 2*/*3; its root-mean-square error across plotted ages is 0.017. At *d* = 400, *N*_*h*_ = 32 and *R* = 169, novel patterns were enrolled prospectively at times 4*R*, 8*R*, 12*R* and 16*R* over eight data seeds, four state seeds and 16 streams, yielding 327 distinct patterns and 1,308 state-specific trajectories. Pointwise 95% intervals use 2,000 hierarchical-bootstrap replicates over data seed, state seed and whole stream. The matched trace intervention uses the same patterns and seeds. Its rate-and-depth-matched Hebbian trace control sets *η*_bin_ and cascade depth equal to those of the anchor for the BTSP-inspired network, and re-optimizes its remaining five parameters at *R* = 31, the Hebbian control’s full-model *R*_0.90_ operating point; it is then traced to memory age 169.

For Fig. 5e, we use the single-level *R* = 169 anchor for the BTSP-inspired network annd hold all remaining parameters and the readout threshold fixed while varying one coupling coefficient. The stationary background is re-solved along the *η*_attn_ sweep; changing *η*_dist_ leaves the dynamics of normalized memory-trace strength unchanged. Both sweeps use *ν*_int_ = 2*/*3 and the full model’s *R* = 169 protocol-weighting novel fraction (*b*_169_ = 0.677108).

### Statistics and reproducibility

Exact chain computations (Figs. 2 and 3) are deterministic; log-domain combinatorics and power iteration (tolerance 10^−14^) solve the exact finite-state chains to the stated numerical tolerance, and the Fig. 2 solver reproduces independently computed reference values digit-for-digit. Held-out accuracies are means over the separate evaluation tensor and 16 model state/event seeds described above; capacities are derived from those per-lag means. Capacity estimates are search-dependent lower envelopes, with adjacent-integer pass/fail localization of each tested crossing. Record-level theory agreement is summarized by mean absolute error, agreement in normalized memory-trace strength by root-mean-square error across plotted ages, and capacity calibration by the geometric-mean reduced-theory/full-model ratio and mean absolute relative error across configurations. Pointwise intervals for memory-trace strength and assignment-opportunity concentration use the hierarchical and crossed-factor bootstrap designs specified above. Model-side random draws and fixed input tensors are generated as described above; for fixed inputs and seeds, discrete state trajectories are bit-exact across software backends. Network-simulation capacity campaigns ran on a single consumer GPU (RTX 4090).

### Data and code availability

All raw study records (tuned parameters, per-lag held-out accuracies, capacity tables), the theory evaluator, and the scripts that regenerate every figure from those records will be deposited in a public repository upon publication.

## Supplementary Information

Plateau-gated one-shot plasticity supports continual recognition memory

These notes provide the mathematical details underlying the main text: exact reductions of the gated jump process (Note 1), the structured-ensemble theory and Gaussian approximation (Note 2), the Curie–Weiss example (Note 3), the stationary single-write reduced theory (Note 4), and a unified symbol dictionary (Note 5). Exact statements are proved or derived explicitly. Approximation steps are identified where they are introduced and assessed against exact finite-state or simulated calculations; the network-specific approximations are catalogued in Supplementary Note 4.

## Supplementary Note 1: exact reductions of the gated jump process

### 1.1 Model and jump-or-stay

Let *w*(*t*) ∈ {±1}^*n*^. In this Note, unstructured inputs are instantiated by independent uniform signed patterns *x*(*t*) ∈ {±1}^*n*^ (independent fair-coin coordinates). The update is equation (1) of the main text, and *s*_u_ = *λ* + *µ*. At a zero local field we retain the preceding synaptic sign; this convention affects only exact equality cases.

#### Lemma 1

(jump-or-stay). *For any scalar gate f* ∈ [0, 1], *away from the tie s*_u_*f* = 1, *w*(*t*+1) = *x*(*t*) *if f* (*w*(*t*), *x*(*t*)) *>* 1*/s*_u_ *and w*(*t*+1) = *w*(*t*) *otherwise. If s*_u_ ≤ 1 *the chain is frozen*.

*Proof*. Coordinate-wise, *w*_*i*_(*t*+1) = sign [1 − *λf*_*t*_]*w*_*i*_ + *µf*_*t*_*x*_*i*_ with *f*_*t*_ = *f* (*w*(*t*), *x*(*t*)). If *x*_*i*_ = *w*_*i*_, the argument is [1 + (*µ* − *λ*)*f*_*t*_]*w*_*i*_; its coefficient is nonnegative and can vanish only at a boundary case covered by the stated tie convention. If *x*_*i*_ = −*w*_*i*_, the argument is [1 − *s*_u_*f*_*t*_]*w*_*i*_, so the coordinate flips exactly when *s*_u_*f*_*t*_ *>* 1. The gate is a scalar, so either all mismatched coordinates flip or none do. For *s*_u_ ≤ 1, *s*_u_*f*_*t*_ ≤ 1 always.

For logistic gates *f* = *σ*_g_(*D/σ*_0_−*θ*_0_) with *σ*_0_ *>* 0, monotonicity of *σ*_g_ converts the event-occurrence condition into *D > κ* with 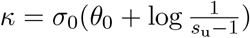, valid in the active regime *s*_u_ *>* 1.

### 1.2 Self-overlap gate

Here *D* = ⟨*w, x*⟩ = *n* − 2*d*_*H*_(*w, x*). Define *R*_*a*_(*κ*) = max({*d* ∈ {0, …, *n*} : *n* − 2*d > κ*} ∪ {0}). Accepted nontrivial updates are precisely those with 1 ≤ *d*_*H*_(*w, x*) ≤ *R*_*a*_; thus *R*_*a*_ = 0 denotes the absence of an accepted state-changing proposal.

#### Theorem 1

(self-overlap gate). *Let s*_u_ *>* 1. *If R*_*a*_ = 0 *the chain is frozen. If R*_*a*_ ≥ 1 *the chain is irreducible and aperiodic on* {±1}^*n*^, *reversible with respect to the uniform distribution, and converges to it. Moreover* E[*w*(*t*) | *w*(0)] = *α*^*t*^*w*(0) *with*

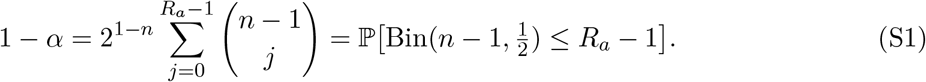

*Proof*. For *y* ≠ *w, P* (*w* → *y*) = 2^−*n*^**1 {**1 ≤ *d*_*H*_(*w, y*) ≤ *R*_*a*_}, which is symmetric in (*w, y*); hence uniform reversibility. One-bit flips have positive probability when *R*_*a*_ ≥ 1 (irreducibility), and the self-proposal *x* = *w* leaves the state unchanged (aperiodicity). A fixed coordinate flips iff the input differs there and the total Hamming distance is at most *R*_*a*_; summing over the *n*−1 other coordinates gives the flip probability 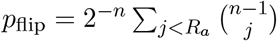 and E[*w*_*i*_(*t*+1) | *w*(*t*)] = (1 − 2*p*_flip_)*w*_*i*_(*t*).

By the binomial large-deviation bound, if 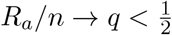 then 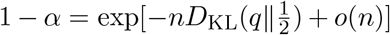: relaxation to the uniform equilibrium is exponentially slow in *n*. This is kinetic freezing: the exponential relaxation time arises from the rarity of accepted updates rather than from an energy barrier (main-text Fig. 2e).

### 1.3 Instructive gate: exact projection onto pathway alignment

Fix the instructive weight vector *v* ∈ {±1}^*n*^ and attenuation 0 *< η* ≤ 1, define the matching-coordinate sets *S* = {*i* : *w*_*i*_*v*_*i*_ = 1} and *T* = {*i* : *x*_*i*_*v*_*i*_ = 1}, and let *m* = |*S*|, *ℓ* = |*T*|, and *u* = |*S*∩*T*| . Here *m* is the pathway-alignment match count and *m/n* is the match fraction.

#### Theorem 2

(exact lumping). *Conditional on m*, 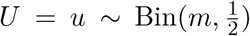 *and* 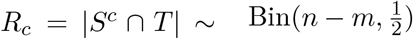 *independently, the proposed next level is ℓ* = *U* + *R*_*c*_, *and the drive equals*

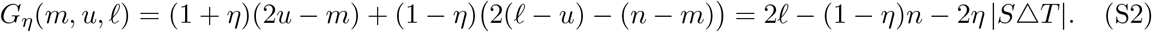

*With the convention that a binomial coefficient vanishes outside its natural range, define*

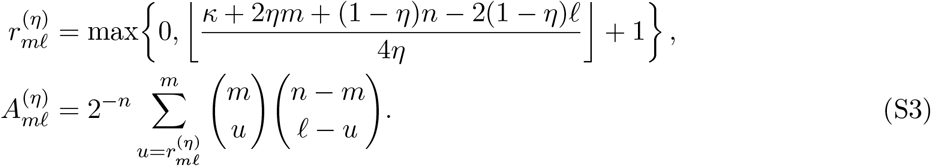

*Consequently m*(*t*) *is a Markov chain with event-occurrence probability* 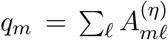 *and held transition matrix Q* = *I* − diag(*q*) + *A*^(*η*)^.

*Proof*. For *i* ∈ *S*, the contribution (*ηw*_*i*_ + *v*_*i*_)*x*_*i*_ equals (1 + *η*)*v*_*i*_*x*_*i*_; for *i* ∉ *S*, it equals (1 − *η*)*v*_*i*_*x*_*i*_. Because *T* = {*i* : *v*_*i*_*x*_*i*_ = 1}, summing gives the first equality in (S2), and *m* + *ℓ* − 2*u* = |*S*△*T*| gives the second. Event occurrence, *G*_*η*_ *> κ*, is monotone in *u* at fixed (*m, ℓ*), which yields the threshold form of *A*^(*η*)^; the binomial counts follow from uniformity of *T* . The transition probability out of *w* depends on *w* only through *m*, which is the strong-lumpability criterion^69^.

#### Theorem 3

(zone structure). *Let s*_u_ *>* 1, 0 *< η* ≤ 1, *and define*

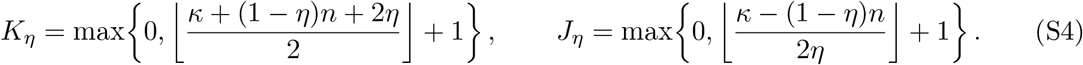

*If κ* ≥ (1+*η*)*n* − 2*η, every state is absorbing. Otherwise, levels m < J*_*η*_ *are frozen; levels J*_*η*_ ≤ *m < K*_*η*_ *are transient, every nontrivial accepted update from them lands in the memory band, and the input x* = *v (probability* 2^−*n*^*) jumps to m* = *n; the memory band* {*m* ≥ *K*_*η*_} *is closed, irreducible and aperiodic, and carries the unique stationary law. For η* = 1, *J*_1_ = max{0, *K*_1_ − 1}; *hence the transient band is empty when K*_1_ = 0 *and otherwise consists only of level K*_1_ − 1.

*Proof*. The largest drive of a nontrivial move is (1 + *η*)*n*− 2*η* (from *m* = *n* − 1 with *x* = *v*), giving the all-frozen threshold. The input *x* = *v* from level *m* has drive (1 − *η*)*n* + 2*ηm*, which exceeds *κ* exactly for *m* ≥ *J*_*η*_; below *J*_*η*_ no input passes. If a nontrivial move lands at *ℓ < K*_*η*_ then, by (S2) with |*S*△*T*| ≥ 1, *G*_*η*_ ≤ 2(*K*_*η*_ − 1) − (1 − *η*)*n* − 2*η* ≤ *κ* by the definition of *K*_*η*_; hence the band is closed and transient levels can only enter it. Irreducibility on the band: from *S* = *m > K*_*η*_ any one-element subset update is accepted (*G*_*η*_ = 2(*m* − 1) (1 − *η*)*n* − 2*η > κ*); from a *K*_*η*_-set any superset update is accepted; and two *K*_*η*_-sets differing by one swap communicate through their union. The self-proposal is accepted on the band, giving aperiodicity.

#### Proposition 1

(single-event entry law). *From a transient level m, entry into the band occurs at a geometric time with success probability* 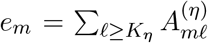, *and the entry level is distributed as* 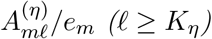.

*Proof*. Until entry the lumped chain cannot move (all accepted updates enter the band), so each step is an independent trial with success probability *e*_*m*_; conditioning on the successful step gives the conditional entry-level distribution.

#### Proposition 2

(stationary law: level symmetry and dwell times). *On the memory band the unique stationary distribution is uniform within each level*, 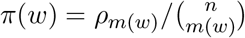, *and the level law factorizes into jump statistics and dwell times: if ξ solves ξ* = *ξB for the embedded accepted-update chain* 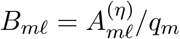.

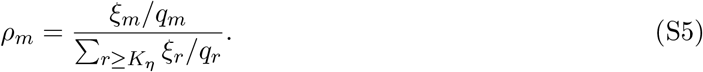

*Proof*. Coordinate permutations preserving *v* are chain automorphisms whose orbits are the levels; uniqueness of the stationary law forces within-level uniformity. From *Q* = *I* − diag(*q*) + *A*^(*η*)^, stationarity *ρ* = *ρQ* is equivalent to *ρ* diag(*q*) = *ρA*^(*η*)^; setting *ξ*_*m*_ ∝ *ρ*_*m*_*q*_*m*_ gives *ξB* = *ξ*, and solving back yields (S5).

Whenever *K*_*η*_ ≤ *n* − 1, irreducibility gives *ρ*_*m*_ *>* 0 for every *m* ∈ {*K*_*η*_, …, *n*}, and therefore E_*ρ*_[*m*] *< n*. Equation (S5) additionally shows how state-dependent dwell times reshape the stationary law; in the high-threshold regimes analyzed in the main text, small occurrence probabilities near the band floor increase the weight of moderately aligned states. The exact stationary accepted-event rate and mean rewrite interval are

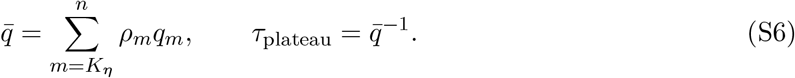

This is the quantity plotted in Fig. 2e. Here a rewrite means the accepted update operation *w x*, including the idempotent operation *x* = *w*. If rewrite is instead required to change the synaptic state, the accepted self-proposal must be removed: 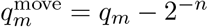 on the memory band.

## Supplementary Note 2: structured input ensembles and the Gaussian fluctuation theory

### 2.1 The gate fixes the topology; the ensemble fixes the flow

Let inputs be drawn i.i.d. from an arbitrary ensemble *ν* on {±1}^*n*^ with mean 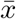, second moment *M*_2_, covariance 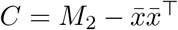, and support *S*. By Lemma 1 the only possible moves are *w* → *y* ∈ *S* passing the gate, with kernel

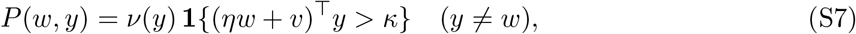

and the self-loop absorbing the remaining probability. The directed graph of allowed moves depends only on (*S, κ, η, v*), not on the positive probability weights {*ν*(*y*) : *y* ∈ *S*}. An arbitrary support may split the graph into several closed communicating classes. For a full-support ensemble, however, the frozen, transient and memory-band geometry is exactly that of Note 1 (Theorem 3); the ensemble probabilities determine the flow and stationary weighting on this fixed graph.

For the self-overlap gate the edge relation is symmetric and the chain is reversible, with stationary law *ν* conditioned on each closed component (*π* = *ν* under full support): the gate changes the kinetics but not the equilibrium. The instructive gate is directed—*w* → *y* requires *η* ⟨*w, y*⟩ + ⟨*v, y*⟩ *> κ* while the reverse edge requires *η* ⟨*w, y*⟩ + ⟨*v, w*⟩ *> κ*—so states similar to *v* are easier to enter than to leave. Under full support, the memory band is irreducible and its stationary law is the unique probability vector satisfying *π* ℒ = 0, where the balance operator is

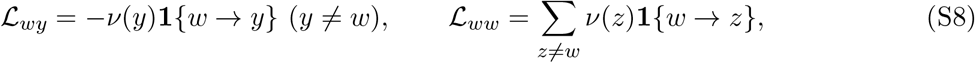

and the held transition matrix is *Q* = *I* − ℒ. The instructive chain generically carries nonzero stationary currents *J*(*x, y*) = *π*(*x*)*ν*(*y*)**1 {***x* → *y*} − *π*(*y*)*ν*(*x*)**1 {***y* → *x*} : a genuinely non-equilibrium steady state. A dwell-time decomposition identical to (S5) holds with *ν*-weighted occurrence probabilities.

### 2.2 Score statistics and the Gaussian closure

All occurrence decisions pass through the scalar drive *S*_*w*_ = *b*(*w*)^⊤^*x* with *b*(*w*) = *ηw* + *v*, whose exact first two moments under *ν* are 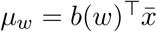 and 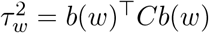. The closure approximates (*x, S*_*w*_) as jointly Gaussian:

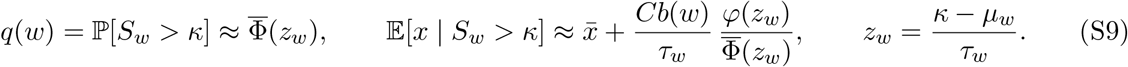

Equation (S9) applies when *τ*_*w*_ *>* 0. If *τ*_*w*_ = 0, the drive is deterministic and occurrence is fixed by whether *µ*_*w*_ *> κ*. The one-step conditional mean of the jump-or-stay update is then 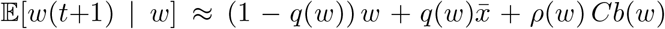 with *ρ*(*w*) = *φ*(*z*_*w*_)*/τ*_*w*_. Two structural consequences: mobility is controlled by the covariance quadratic 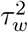, and the instructive weight vector acts only as *Cv*—filtered through the input covariance.

### 2.3 Order parameters, drift closure, and the self-consistent mean

In the eigenbasis *Cu*_*r*_ = *λ*_*r*_*u*_*r*_ write 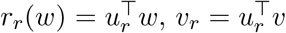. For balanced ensembles 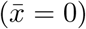 the closure gives the modal drift

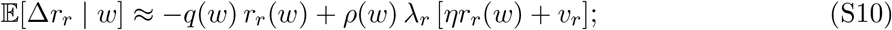

modes couple only through the global scalars *q, ρ*. For the covariance-filtered alignments 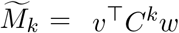,

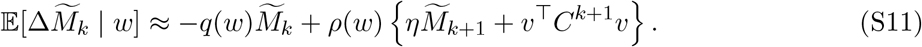

This hierarchy is generally infinite; it becomes finite after reduction by the distinct eigenvalues (equivalently, the minimal polynomial) of *C*. In particular, 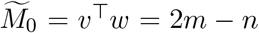 fixes the exact support, whereas 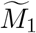 determines its drift.

Deriving a stationary mean requires a second, representative-state closure in addition to Gaussianizing the drive. Specifically, set 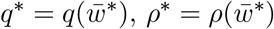 and approximate

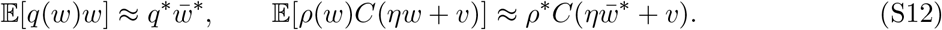

The zero-drift condition then yields the self-consistent mean in main-text equation (4). With 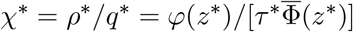 and *z*^∗^ = *κ/τ*^∗^,

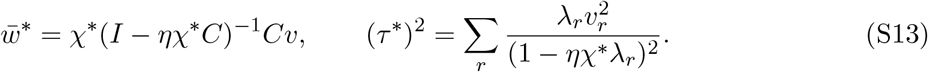

The positive-gap condition 1 − *ηχ*^∗^*λ*_*r*_ *>* 0 supplies invertibility and frozen-coefficient linear stability. The binary bounds 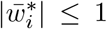 and exact band floor 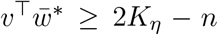 are additional necessary admissibility diagnostics, but they are not sufficient to establish a binary stationary law with the proposed mean. Where they fail, the Gaussian mean is only a formal local branch. Such sectors are identified explicitly in Supplementary Note 3. Every branch in the two aligned sweeps of main-text Fig. 3c satisfies these constraints.

### 2.4 Linear-noise covariance

Let *m*_1_(*w*) = E_*ν*_ [*x***1 {***S*_*w*_ *> κ*}], *G*(*w*) = E_*ν*_ [*xx*^⊤^**1{***S*_*w*_ *> κ*}], and *D*(*w*) = *m*_1_(*w*) − *q*(*w*)*w*. The exact one-step conditional moments of the jump-or-stay process are

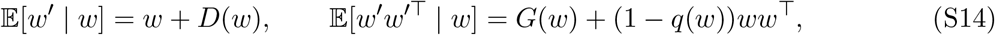

so the exact conditional innovation covariance is

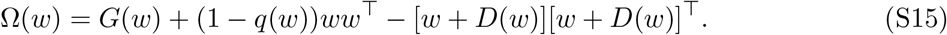

For a balanced joint-Gaussian proposal, let *c*(*w*) = *Cb*(*w*)*/τ*_*w*_. The corresponding truncated second moment is

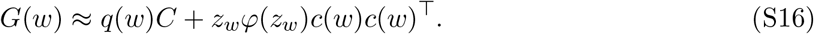

Equations (S15) and (S16) expose the stay-mixture and truncation terms entering a full local covariance calculation.

For the closed-form modal diagnostic used here, two further linear-noise approximations are made: derivatives of *q* and *ρ* are omitted from the drift Jacobian, and the rank-one truncation and stay-mixture corrections in Ω are dropped. Thus *J*_0_ = −*q*^∗^(*I* − *ηχ*^∗^*C*) and Ω_0_ = *q*^∗^*C*. The discrete Lyapunov equation Σ^∗^ = (*I*+*J*_0_)Σ^∗^(*I*+*J*_0_)^⊤^ + Ω_0_ is diagonal in the eigenbasis of *C*:

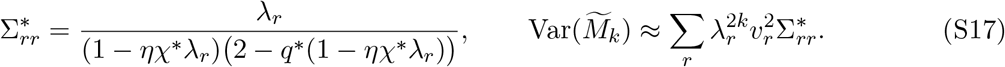

Growth of a modal variance as *ηχ*^∗^*λ*_*r*_ → 1 signals loss of validity of the frozen-coefficient approximation; the exact binary chain has bounded covariance. Accordingly, equation (S17) is a linear-noise approximation, not an exact stationary covariance.

## Supplementary Note 3: the Curie–Weiss worked example

### 3.1 Ensemble, covariance, and criticality

The structured-input case is instantiated here by the correlated Curie–Weiss family 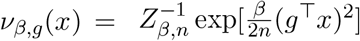, balanced and full-support for every finite *β*. With *S* = *g*^⊤^*x*,

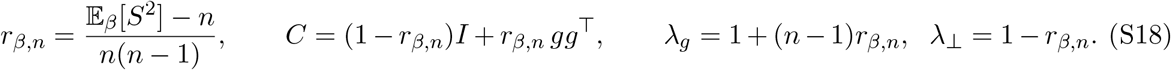

Because *λ*_*g*_ = E_*β*_[*S*^2^]*/n*, it is *O*(1) for *β <* 1 and *O*(*n*) for *β >* 1 in the thermodynamic limit. In the latter regime, *S/n* develops modes at ±*m*_*β*_, where *m*_*β*_ = tanh(*βm*_*β*_). Thus *β* = 1 is the thermodynamic-limit critical point; at finite *n*, including the value used here, the change is a smooth crossover toward the two directions ±*g* (main-text Fig. 3b).

### 3.2 Exact two-parameter projection

Split coordinates into *I*_+_ = {*i* : *g*_*i*_ = *v*_*i*_} (size *N*_+_ = (*n* + *h*)*/*2) and *I*_−_ (size *N*_−_ = (*n* − *h*)*/*2), where *h* = *g*^⊤^*v* and *n*±*h* are even. Let *a*_+_(*w*), *a*_−_(*w*) count matches to *g* inside *I*_+_, *I*_−_. Signed permutations fixing *g* and *v* act transitively on each (*a*_+_, *a*_−_) set, and both the ensemble and the gate are invariant, so the chain lumps exactly onto (*a*_+_, *a*_−_), equivalently onto (*G, V* ) = (2(*a*_+_+*a*_−_) − *n*, 2(*a*_+_ *a*_−_) *h*). Thus *G/n* is the structure overlap and *V/n* is the signed instructive overlap, while the instructive match count *a*_+_ + *N*_−_ − *a*_−_ plays the role of the alignment *m*. From orbit (*a*_+_, *a*_−_) to orbit (*α*_+_, *α*_−_) the accepted flux is

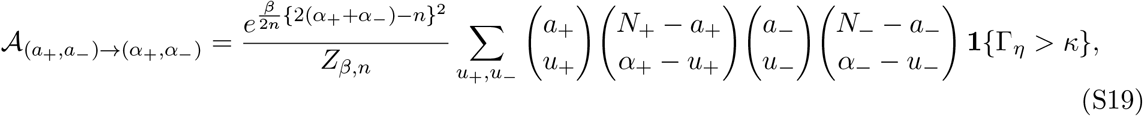

Γ_*η*_ = *η*{*n* − 2(*a*_+_+*a*_−_+*α*_+_+*α*_−_) + 4(*u*_+_ + *u*_−_)} + 2(*α*_+_ − *α*_−_) − *h*, restricted to the active orbits *a*_+_ + *N*_−_ − *a*_−_ ≥ *K*_*η*_; the sums run over all integers for which the binomial coefficients are defined. For distinct orbits *ω, ω*^*′*^, set *Q*_*ωω*_*′* = *A*_*ω*→*ω*_*′* and *Q*_*ωω*_ = 1 − ∑ _*ω′ ω*_ *A*_*ω*→*ω*_*′* . The exact stationary orbit law solves *πQ* = *π* (equivalently, *π*(*I* − *Q*) = 0) and was computed by dwell-time decomposition and power iteration, without sampling.

### 3.3 Closed-form resolvent and assessment

Because *C* has two eigenvalues, (S13) closes on two scalars. With modal gaps Δ_*g*_ = 1 − *ηχ*^∗^*λ*_*g*_, Δ_⊥_ = 1 − *ηχ*^∗^*λ*_⊥_:

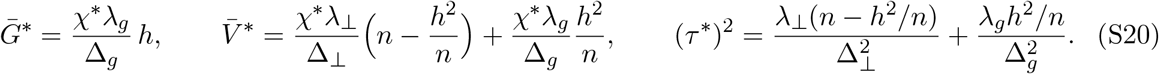

Admissibility uses 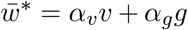 with *α* = *χ*^∗^*λ /*Δ, *α* = (*h/n*)(*χ*^∗^*λ /*Δ ^−^ *χ*^∗^*λ /*Δ ): binary bounds |*α*_*v*_ ± *α*_*g*_| ≤ 1 and the band floor on 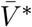. On the sweep of main-text Fig. 3c (*n* = 64, *h/n* = 0.5, *η* = 0.5; *κ* = 0 and 20), for *β* ≤ 1 the closure predicts the exact means to r.m.s. error 0.006 (*G/n*) and 0.005 (*V/n*) at *κ* = 0, and 0.048/0.032 at *κ* = 20. For *β >* 1 errors grow in the dominant-pattern overlap (*G/n* r.m.s. 0.15 at *κ* = 0). For an orthogonal instructive vector, the stable single-centered branch also disappears over part of the supercritical, high-threshold plane (dashed boundary in Supplementary Fig. S1c; gray region in Supplementary Fig. S1a)—a failure mode predicted by the closure’s own diagnostics (*λ*_*g*_ macroscopic, ensemble bimodal, Δ_*g*_ → 0).

The coefficient maps in main-text Fig. 3d display the continuous formal positive-gap resolvent branch to expose its parameter dependence. In the high-*β*, high-*κ* corner this branch violates the binary bounds |*α*_*v*_ ± *α*_*g*_| ≤ 1 and therefore does not define an admissible stationary mean of the binary chain. That sector indicates saturation of the single-well closure and requires a constrained or multimodal extension.

**Figure S1:**
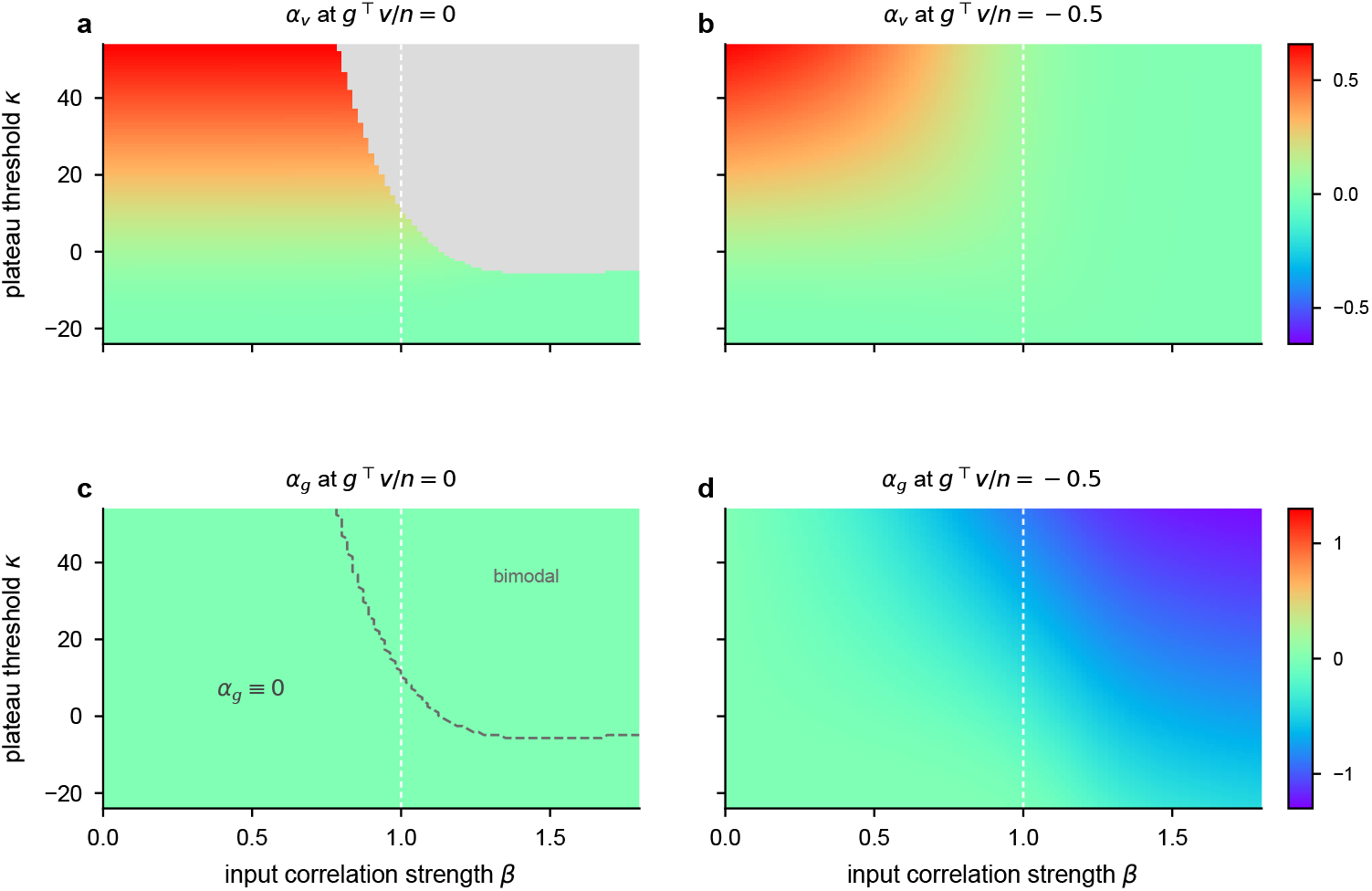
The Gaussian resolvent across instructive–structure overlaps. Formal positive-gap coefficients in 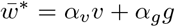 across the (*β, κ*) plane (*n* = 64, *η* = 0.5) for an orthogonal instructive vector (*g*^⊤^*v/n* = 0; **a**,**c**) and an anti-aligned instructive vector (*g*^⊤^*v/n* = − 0.5; **b**,**d**), complementing the aligned case in Fig. 3d. **a**,**b**, The instructive coefficient *α*_*v*_ is even in the instructive–structure overlap; the anti-aligned plane therefore coincides with the aligned plane. **c**,**d**, The structure coefficient *α*_*g*_ is odd in the overlap: it vanishes for an orthogonal instructive vector and reverses sign under an anti-aligned instructive vector. Gray in **a** denotes the absence of a positive-gap single-well solution; the dashed curve in **c** marks the same boundary. In **c**, *α*_*g*_ = 0 is imposed by the exact *G* ↦ − *G* symmetry, including beyond that boundary. Colored formal branches can still violate the binary or memory-band diagnostics described in Supplementary Note 3.

#### The orthogonal plane

At *h* = 0 the coordinate classes *I*_*±*_ have equal size *n/*2, and any signed permutation exchanging them leaves the proposal (even in *g*) and the gate (blind to *g*) invariant while reversing the sign of *G*: the exact stationary law is symmetric under *G* ↦ − *G*, so the mean structure component vanishes identically, ⟨*G*⟩ ≡ 0, on the whole (*β, κ*) plane (verified to machine precision by the orbit chain, including deep in the supercritical high-threshold regime, e.g. |E[*G*]| *<* 10^−13^ at *κ* = 50, *β* = 1.5). The single-centered Gaussian loses its stable positive-gap branch there (*ηχ*^∗^*λ*_*g*_ → 1) as the symmetric law becomes bimodal in the ±*g* sectors. The exact symmetry motivates a two-component ansatz with equal weights and centers *α*_*v*_*v* ±*ag*: the mixture retains the instructive component and cancels the structure component. Determining the amplitude *a* and the within-component covariance requires an additional nonlinear saturation closure, which is not developed here. Supplementary Fig. S1c therefore reports *α*_*g*_ = 0 across the whole *h* = 0 plane and marks the single-well stability boundary (dashed); the instructive component in that regime is not symmetry-protected and is left blank in Supplementary Fig. S1a.

## Supplementary Note 4: reduced theory for a stationary single write

This Note develops a large-network reduced theory of the BTSP-inspired continual-recognition network. It isolates a stationary single-write experiment: a pattern is assigned to one memory unit and written once, independent novel patterns subsequently interfere with that unit, and familiar and novel probes are read without further plasticity. This controlled task separates three questions that are entangled in the full model: which memory unit is selected, how its synaptic engram and matched no-write control evolve, and how their causal engram contrast affects a population decision. Conditional on the supplied network parameters and readout threshold, the reduction introduces no additional dynamical or moment-closure coefficient fitted to recognition accuracy. Its dependency chain is

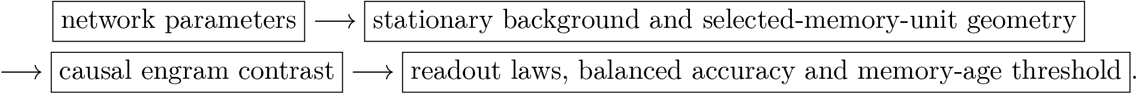

The five subsections below correspond to Sections 1–5 of the standalone theory report. They contain the theoretical construction only; numerical tests and parameter-specific results are excluded.

### 4.1 Network, task and approximation framework

#### Two computational pathways

An input is a binary vector *x* ∈ {0, 1}^*d*^ with exactly *k* active coordinates,

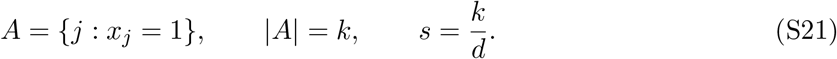

The memory layer contains *N*_*h*_ memory units, each corresponding to one row. Plastic synapses *S*_*ij*_ ∈ {−1, +1} change with experience, whereas *W*_*ij*_ ∈ {− 1, +1} are fixed instructive weights. Thus rows *S*_*i*_ and *W*_*i*_ play the roles of the single-neuron plastic and instructive vectors *w* and *v*, respectively. Unlike the dense signed inputs in Supplementary Notes 1–3, this reduced theory uses exact-*k* sparse binary inputs in *d* dimensions. It takes each instructive row to contain *d/*2 entries of each sign. This balance enforces zero row mean and, relative to centering i.i.d. binary rows, removes centered-norm heterogeneity. It is an explicit idealization of the network. For row *i*, define

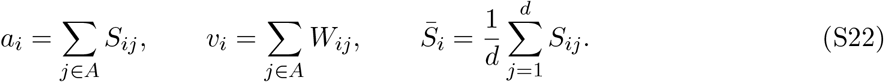

The credit-assignment score and its normalized softmax are

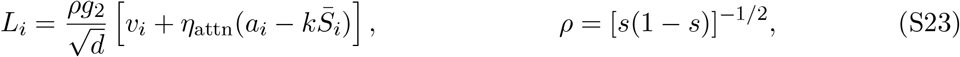

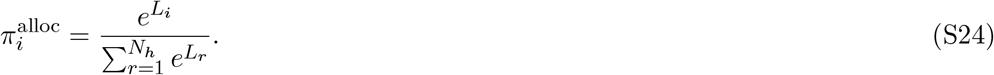

Thus *g*_2_ is the assignment gain, while the proximal assignment coupling *η*_attn_ sets the contribution of the current plastic state to the credit-assignment score alongside the fixed instructive drive. Because the softmax is normalized, every presentation carries unit credit-assignment mass: this reduced theory has no additional plateau threshold, eligibility rejection or no-write branch. Accordingly, *g*_2_ is not a network analogue of the single-neuron plateau threshold *κ*. In the full model, this normalized update assignment remains active on every presentation while familiarity-dependent plasticity suppression separately scales the plasticity passes.

Recognition uses the same two drives through a distinct coupling,

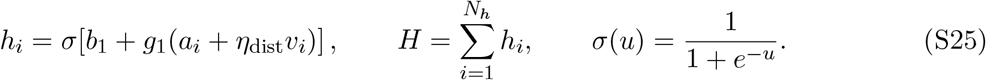

Here *h*_*i*_ is a memory-unit response, *H* is the population response, and the full model maps *H* to its continuous familiarity score *y*. For *w*_2_ *>* 0, the thresholded decision reports a probe as familiar when

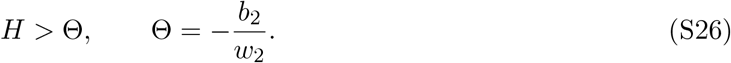

Only the ratio −*b*_2_*/w*_2_ therefore enters the hard decision. In biological terms, the stable instructive pathway and the current plastic state jointly determine the selected memory unit for an update, whereas the distal readout coupling in equation (S25) determines how the plastic synaptic state is expressed during recognition. Credit assignment and memory recall can therefore be coupled without being identical computations.

#### Stationary single-write task

Each episode begins by drawing a balanced *W* and then drawing *S*, conditional on *W*, from the stationary product law derived in Supplementary Note 4.2. A tagged exact-*k* pattern is then drawn, assigned to one memory unit, and written once. It is followed by *a* − 1 independent novel distractors. Finally, a familiar probe containing the tag and an independently drawn novel probe are presented to the same terminal state with plasticity disabled. Balanced accuracy is computed using the one fixed readout threshold in equation (S26). Accordingly, *a* = 1 means immediate post-write readout. The background is stationary and the tag is never deliberately replayed, so the calculation depends on its age *a*, not on an additional dataset horizon. This experiment is a reduced theory of the full model, rather than an identity for the refreshing single-repeat protocol with familiarity-dependent plasticity suppression. The reduced theory then uses conditional occupancies, Gaussian credit-assignment scores and categorical memory-unit selection.

The full model’s exact softmax distributes normalized credit-assignment weights over memory units. The reduced theory preserves those fractional assignment weights in equation (S24), then samples one selected memory unit,

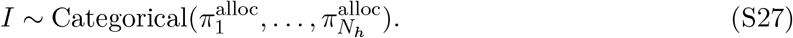

This preserves the marginal assignment probability 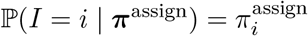, but replaces within-presentation cross-row correlations by exclusive selection. It becomes most accurate as the optimized softmax approaches hard selection. The selected memory unit undergoes the collapsed local cycle

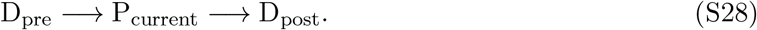

Potentiation targets the current active set and flips an eligible negative synapse with probability *η*_bin_. Each depression pass uses an independent random exact-*k* set and flips a targeted positive synapse with probability

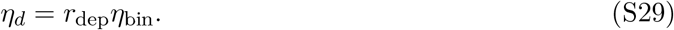

Here *r*_dep_ is the depression-to-potentiation ratio; in the studied BTSP-inspired network, *η*_*d*_ = *η*_dep_ and *r*_dep_ = 1*/*2. This local cycle retains the competition between one-shot writing and homeostatic depression while omitting dependencies on adjacent presentations.

#### Approximation contract

Within the declared reduction, exact-*k* finite-population coefficients, the conditional two-state synaptic kernel and the finite tagged-row count lattices are retained explicitly. The principal closures are the balanced binary instructive rows, conditional independence within the two instructive-sign classes, independent Gaussian row scores, coefficient-free approximations to two selected-score moments, a second-order Hermite projection from memory-unit selection to coordinate enrichment, categorical selection, a stationary reservoir unaffected by one tagged memory unit, exchangeable collisions, conditionally independent tagged coordinates and a two-moment nonnegative population remainder. These assumptions define where the theory is exact conditional on supplied quantities and where it is a large-network approximation; none introduces a coefficient fitted to balanced accuracy. Relative to the full-model simulations, the reduced theory additionally uses single-level binary synapses, exactly balanced instructive rows, the centered plastic credit-assignment drive in equation (S23), independent depression sets, disabled familiarity-dependent plasticity suppression and no tag replay. Full-model simulations can instead use metaplastic cascades, uncentered plastic credit assignment, distributed softmax credit assignment and dependencies between adjacent presentations. Unless otherwise stated, the construction assumes even *d >* 2, 0 *< k < d, N*_*h*_ *>* 1, 0 *< η*_bin_ ≤ 1, 0 ≤ *η*_*d*_ ≤ 1, *g*_2_ ≥ 0, valid derived flip probabilities, *V*_*L*_ *>* 0, and *U*_*sw*_ + *D*_*sw*_ *>* 0 for every retained instructive-sign class. The Hebbian-control extension additionally requires *η*_*ℓ*1_ ≤ 1, and its hard-selection and increasing-response formulas assume *g*_2_ *>* 0 and *g*_1_ *>* 0, respectively.

### 4.2 Stationary self-consistency and selected-memory-unit geometry

#### Two background coordinates

For balanced instructive rows, a synapse belongs to one of two exchangeable instructive-sign classes. Their stationary positive probabilities are

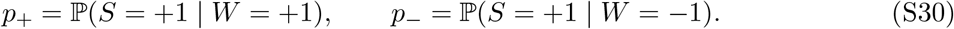

The conditional product closure is conveniently summarized by

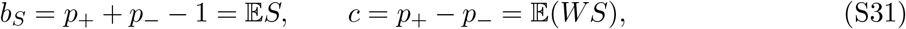

where *b*_*S*_ is the plastic bias and *c* is the plastic–instructive correlation. Both are required: correlation changes which memory units receive updates and hence changes the transition currents that maintain the background.

Uniform exact-*k* sampling gives the finite-population factor

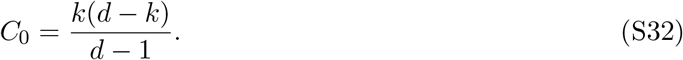

Because *k* = *sd*, the centered assignment drive has the exact representation

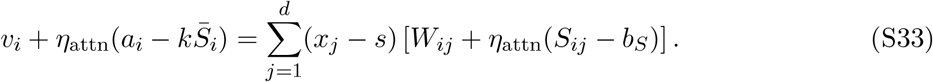

Under the conditional product law, the coordinate contribution *Y* = *W* + *η*_attn_(*S* − *b*_*S*_) has variance

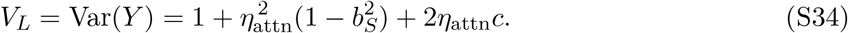

The Gaussian score closure replaces each row’s empirical coordinate variance by this product-law value. Its expectation differs from *V*_*L*_ by *O*(*d*^−1^), while fixed-realization row-to-row fluctuations are *O*_P_(*d*^−1*/*2^). Approximating the row logits by *L*_*i*_ ≃ *σ*_*L*_*ζ*_*i*_, with independent *ζ*_*i*_ ∼ *N* (0, 1), then defines the dimensionless assignment-logit scale

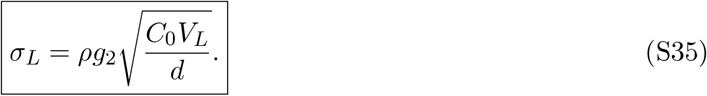

#### Selected-score moments

Let 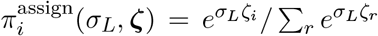, and let *φ* and φ denote the standard-normal density and cumulative distribution. The first two Hermite moments of the score seen by the selected memory unit are

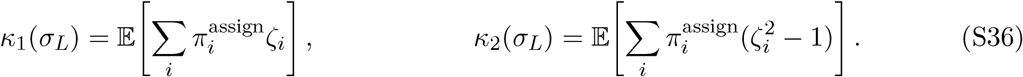

At zero gain selection is uniform; at infinite gain the selected memory unit has the maximum of *N*_*h*_ Gaussian scores. Consequently,

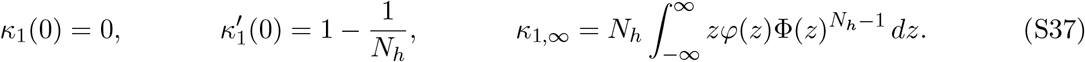

The coefficient-free interpolation

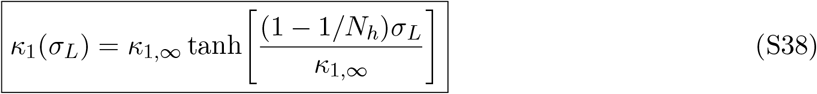

has the exact value and slope at the origin and the exact hard-selection endpoint under the Gaussian score model. Gaussian integration by parts similarly gives

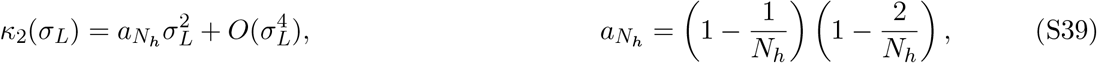

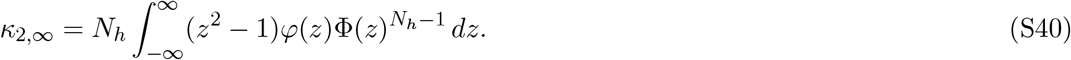

For *N*_*h*_ *>* 2, writing 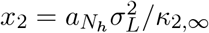, the minimal root–Padé closure is

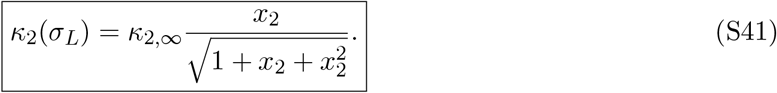

It has the exact leading small-gain coefficient and hard-selection endpoint. For *N*_*h*_ = 2, *x*_2_ is not formed: both limiting quantities vanish and *κ*_2_ ≡ 0.

#### From memory-unit selection to synaptic enrichment

For instructive sign *s*_*W*_ ∈ {−1, +1} and current plastic sign *s*_*S*_ ∈ {−1, +1}, define

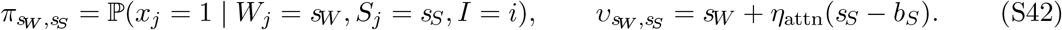

A second-order Hermite projection of coordinate activity onto the standardized row score gives

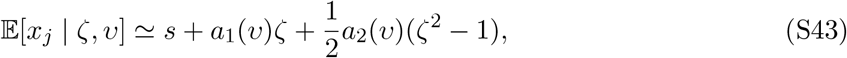

with the exact-*k* coefficients

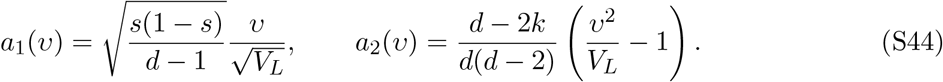

Both coefficients take the row’s centered coordinate scores *W*_*ij*_ + *η*_attn_(*S*_*ij*_ *− b*_*S*_) to sum to zero over *j* and to have squared norm *dV*_*L*_, as in equation (S35); this neglects the residual plastic row sum 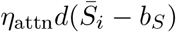, a relative *O*_P_ (*d*^−1*/*2^) correction of the same order as the row-to-row fluctuations above. Averaging over the selected-score law yields four enrichment probabilities,

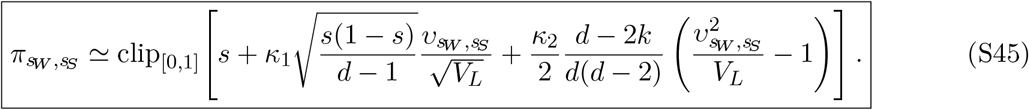

The quadratic term contains the interaction 2*η*_attn_*s*_*W*_ *s*_*S*_. It restores the leading instructive-by-state selection effect omitted by first-order regression, scales as *d*^−1^ at fixed sparsity, and vanishes at zero assignment gain or when *η*_attn_ = 0. On an interior branch the projection respects the exact-*k* mean constraint. Componentwise clipping is a bounded closure and can violate that constraint slightly; no exact normalization claim is made for a clipped branch.

#### Plastic transition currents and fixed point

A coordinate is included and successfully depressed in either random depression pass with probability

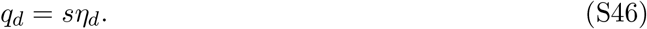

In one selected-memory-unit plasticity cycle, the class-*s*_*W*_ probabilities of an upward (−1 → +1) and downward (+1 → −1) transition are

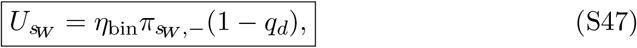

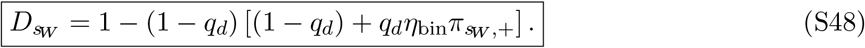

Here 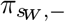 and 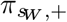 abbreviate 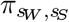 at *s*_*S*_ = − 1 and +1, respectively. The second expression includes repair by potentiation when pre-depression and the current active set meet on the same synapse. Stationary upward and downward currents balance when

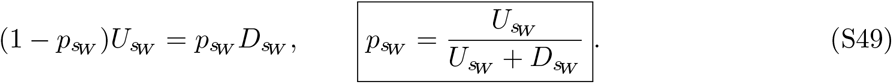

This is a two-dimensional self-consistency problem because the rates depend on the selected-memory-unit geometry generated by *p*_+_ and *p*_−_. One iteration has the explicit dependency order

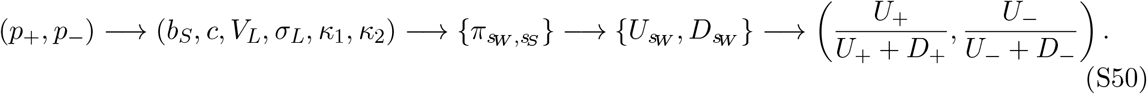

A damped bounded fixed-point or residual solve selects the stationary pair. Because clipping makes the map non-smooth, no general uniqueness claim is made: the initialization, selected branch and final residual are part of the solver contract. The explicit selection-frequency factor 1*/N*_*h*_ cancels between the two currents and therefore controls their common time scale, not their ratio. Width can nevertheless change the equilibrium indirectly through the *N*_*h*_-dependent selected-score moments.

Once the fixed point is known, the active coordinates of the selected memory unit have instructive-sign mass

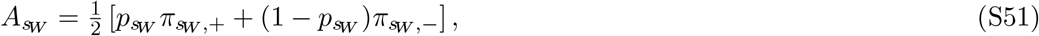

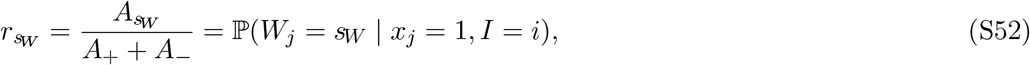

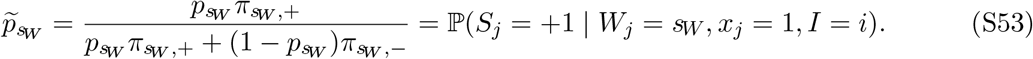

Thus 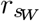 describes which instructive signs occur on the active coordinates of the selected memory unit, and 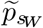 describes their positive plastic fraction before the write. Together with 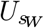 and 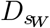, these quantities close the tagged synaptic engram dynamics.

### 4.3 Synaptic engram initialization and collision-limited evolution

#### Matched no-write control

Memory-unit selection enriches the tagged active set before any synapse is changed. Comparing the selected memory unit after writing with an unselected stationary memory unit would therefore confound this pre-existing selection bias with learning. We instead introduce a matched no-write control copy of the selected memory unit, expose it to the same later interference, and omit only the tagged plasticity cycle. Its positive probability in instructive-sign class *s*_*W*_ at immediate readout is

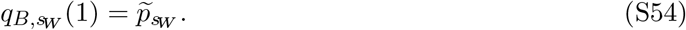

This paired counterfactual makes the causal engram contrast vanish identically when potentiation is removed.

#### Immediate one-shot write

For the tagged pattern, every active coordinate is eligible for potentiation rather than being sampled with the selection-averaged probability 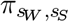. Conditional on the pre-write plastic sign, the probability of being positive after the complete depression– potentiation–depression cycle is

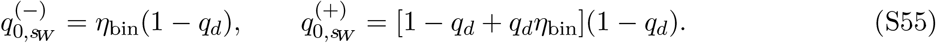

Mixing over the selected pre-write composition gives

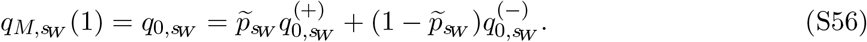

The subscripts *M* and *B* denote the memory and its matched no-write control; *a* = 1 is the common immediate-readout convention.

#### Collision dynamics

When the tagged memory unit is later selected for a distractor, each tagged coordinate follows the same two-state kernel as the stationary background. For either member of the matched pair,

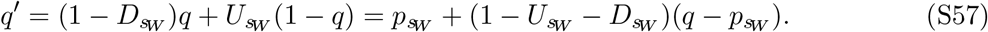

Define 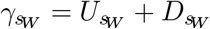 . The class probability after *n* collisions with the selected memory unit is

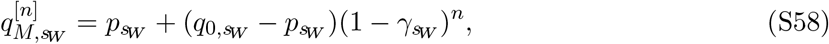

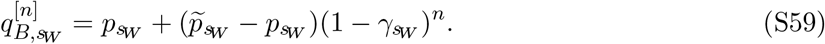

The tagged memory unit undergoes a plasticity cycle at every collision and can change microscopically, but it contributes only *O*(1*/N*_*h*_) to the reservoir. The stationary-reservoir closure therefore keeps 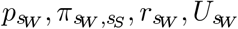 and 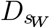 fixed instead of re-solving the population background at every age.

Let *ν*_int_ be the fraction of chronological presentations treated as effective novel interference. The all-fresh task has *ν*_int_ = 1; the zero-order surrogate for the single-repeat stream uses its nominal novel fraction, *ν*_int_ = 2*/*3. More generally, a mixed stream replaces the fresh-update kernel *K* by (1 − *ν*_int_)Id + *ν*_int_*K*. This state-independent thinning multiplies both stationary currents by the same factor and hence changes trace retention, but neither *p*_*sW*_ nor the immediate write. Exchangeability gives marginal selection probability 1*/N*_*h*_. The resulting collision rate is *ν*_int_*/N*_*h*_; the following count model additionally treats selection indicators and thinning decisions as independent and state independent across presentations. Under this closure,

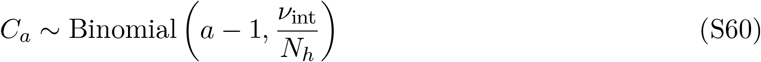

is the number of effective collisions before a probe. Averaging over this collision count gives the exact finite-binomial survival factor and its Poisson limit,

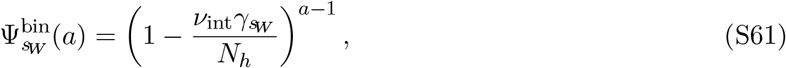

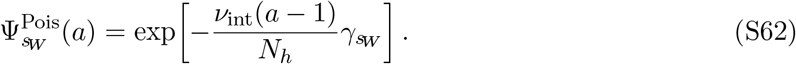

With either declared choice of 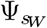,

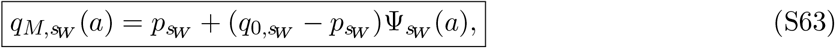

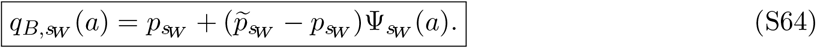

The finite-binomial law is retained when the full tagged-memory-unit mixture is needed. The Poisson form makes the scaling transparent and applies, for example, as *N*_*h*_ → ∞ with (*a* − 1)*/N*_*h*_ = *O*(1); more generally its logarithmic error is small when 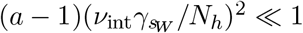. Treating a familiar fraction 1 − *ν*_int_ as read-only is a protocol approximation, not a fitted correction: it omits residual effects of familiarity-dependent plasticity suppression and history-dependent assignment on those events.

#### Causal engram contrast and trace lifetime

The causal class contrast is

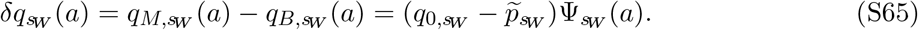

Because a change in positive probability changes the mean of a binary {−1, +1} synapse by twice that amount, the causal engram contrast is

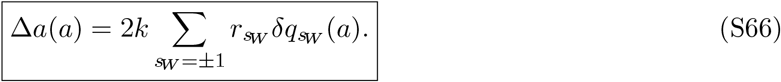

The two instructive-sign classes need not decay at the same rate. When the initial causal contrast is nonzero and *ν*_int_*γ*_eff_ *>* 0, a compact lifetime is obtained from the initial-contrast-weighted contraction rate,

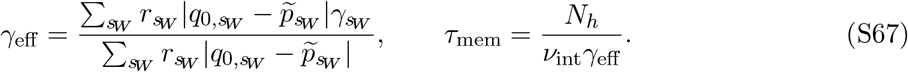

This absolute-amplitude summary need not equal the initial decay rate of the signed aggregate if the two class contrasts have opposite signs. It exposes the principal protection afforded by sparse credit assignment: widening the memory layer lowers the frequency with which any particular memory unit is selected, while the potentiation and depression probabilities jointly set both the initial causal engram contrast and its loss per collision.

### 4.4 Readout, balanced accuracy and memory-age threshold

#### Finite single-memory-unit laws

Recognition requires three response distributions. *P*_gen_(*h*) is the response of a generic stationary memory unit to a novel input, *P*_*M*_ (*h*; *a*) is the response of the written tagged memory unit to its familiar input, and *P*_*B*_(*h*; *a*) is the response of the matched no-write control for the selected memory unit to that same input. The difference between *P*_*M*_ and *P*_*B*_ carries the causal memory; *P*_gen_ supplies the remaining population background.

For a generic memory unit, let *ℓ* be the number of active input coordinates with *W* = +1, and let *Z*_+_ and *Z*_−_ count positive plastic synapses in the two instructive-sign classes. Then

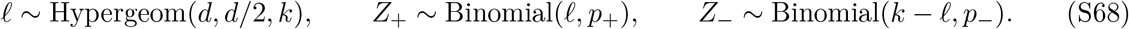

Every finite count triple maps to plastic and instructive drive components, their combined pre-sigmoid drive, and a memory-unit response,

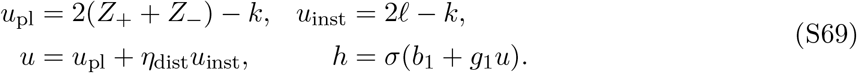

Summing the resulting product-law probabilities gives *P*_gen_. For a tagged memory unit, the composition-independence closure takes *ℓ* ∼ Binomial(*k, r*_+_); selection conditioning induces finite-coordinate dependencies that this marginal law does not retain. Conditional on the collision count in equation (S60), the positive probabilities are 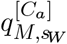 or 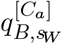. Summing over *C*_*a*_ and the two conditional sign counts gives *P*_*M*_ and *P*_*B*_. Keeping these lattices matters because applying a steep sigmoid to a small integer count is not equivalent to applying it only to the count mean. For the tagged selected memory unit, *Z* = *Z*_+_ + *Z*_−_ is its synaptic engram, and E[*Z*]*/k* is the normalized memory-trace strength.

The nonlinear memory-unit response contrast and the first two population moments are

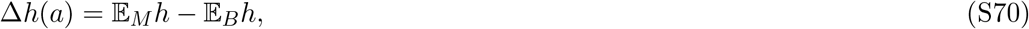

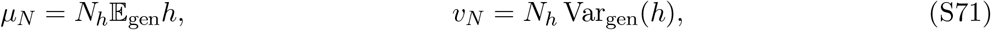

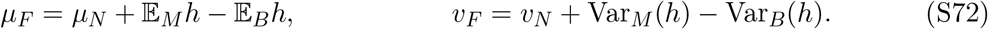

#### A shared nonnegative population remainder

Sums of sparse sigmoid responses are nonnegative and generally skewed. For *µ >* 0 and *v >* 0, let *F*_Γ_(*x*; *µ, v*) denote the CDF of a gamma law with mean *µ* and variance *v*, that is, with shape *µ*^2^*/v* and scale *v/µ*. Define the formal remainder moments

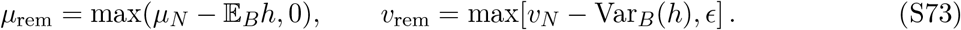

*ϵ >* 0 is a fixed numerical variance floor used only to keep the gamma parameterization well defined. If *µ*_rem_ = 0, the remainder is defined as a point mass at zero rather than by the singular gamma parameterization. When either maximum in equation (S73) is active, nonnegativity is preserved but exact moment matching is necessarily relaxed.

The same formal remainder is used for both hypotheses,

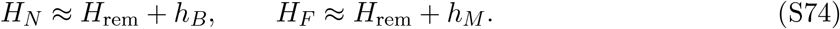

This is two-moment bookkeeping, not a literal decomposition into one selected memory unit and *N*_*h*_ − 1 independent generic memory units. In particular, its rematching can give the approximated novel CDF a weak memory-age dependence even though the stationary generic background is age independent. Sharing the remainder is nevertheless essential: if *P*_*M*_ = *P*_*B*_, the familiar and novel laws coincide and the closure cannot manufacture residual memory. Their population CDFs are the finite convolutions

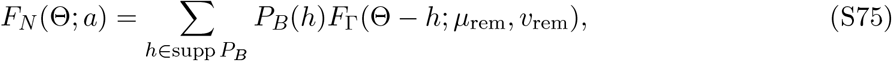

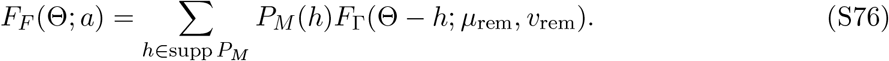

With the familiar-if-high convention,

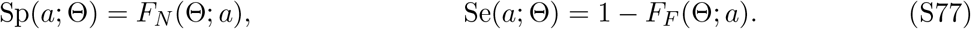

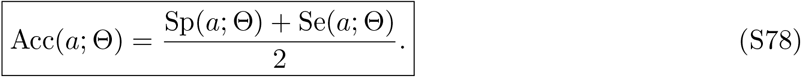

The readout threshold Θ = −*b*_2_*/w*_2_ is supplied by the deployed network, or may be chosen once at a declared design age and then held fixed. Retuning it independently at every age would define a sequence of classifiers rather than the retention range of one classifier. For a target *α* ∈ (1*/*2, 1), provided the passing set is nonempty and bounded,

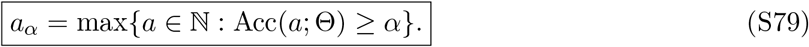

The finite-lattice calculation determines this crossing and should verify both sides rather than assume monotonicity. A useful linearized design relation, valid for a positive decaying contrast with Δ*h*_0_ *>* Δ*h*_req_ *>* 0, is

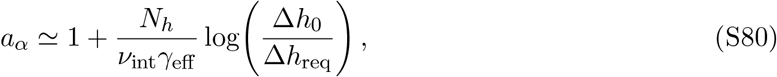

where Δ*h*_0_ is the initial memory-unit response contrast and Δ*h*_req_ is the contrast required at the decision tail. This approximation separates trace lifetime from readout demand; it is a scaling guide, not a replacement for the finite distributions.

#### How the two coupling coefficients act

The two coupling coefficients enter at different stages,

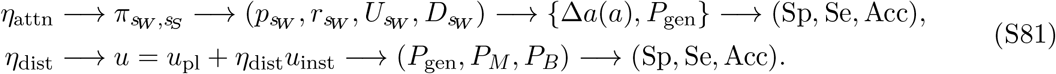

Proximal assignment coupling therefore changes both the memory-bearing reservoir and the novel background. Distal readout coupling instead reads out an already fixed reservoir. Intermediate coupling implements partial, rather than complete, separation of assignment from storage.

In the saturated-selection regime, *κ*_1_ and *κ*_2_ have reached their hard-selection limits, so changing *η*_attn_ primarily changes which coordinates are selected rather than the concentration of credit assignment. Let a prime denote a total derivative with respect to *η*_attn_ after re-solving the stationary pair. Then

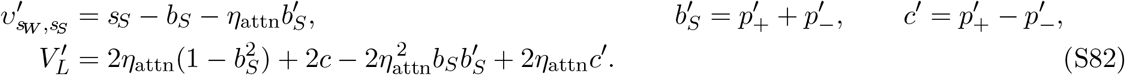

For an unclipped enrichment probability,

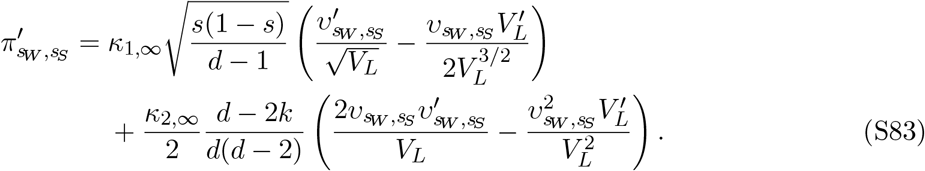

The derivative is zero along a probability-clipped branch and is non-smooth at its boundary. Away from saturation, the corresponding 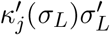 terms must also be retained. The induced transition-probability and fixed-point derivatives are

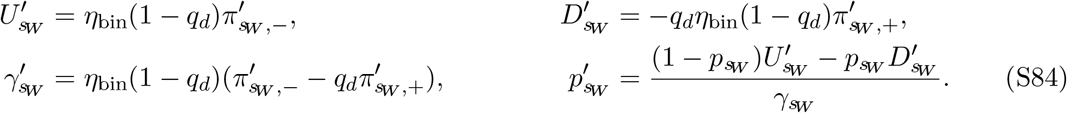

Together with equation (S83), the final relation is a pair of linear equations for 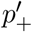 and 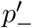. On a state-enriching branch where 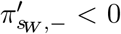 and 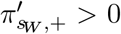, selected-memory-unit plasticity cycles become less likely to change a synapse in the selected memory unit, and retention improves. This sign pattern is a regime property, not a universal theorem.

On an interior branch with 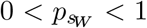 and 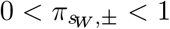, the same selection can raise the positive pre-write fraction,

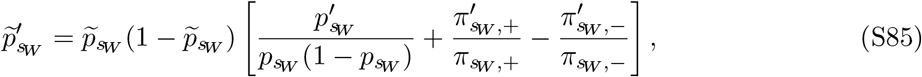

leaving less room for a potentiating write. Indeed,

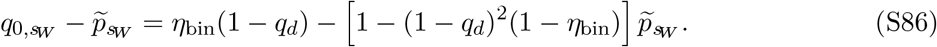

When the class contrast is positive, the Poisson survival form makes the competition explicit,

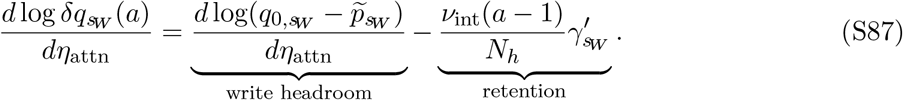

The write cost acts immediately, whereas the benefit of slower decay grows with memory age. credit assignment also changes classification through the novel-memory-unit drive,

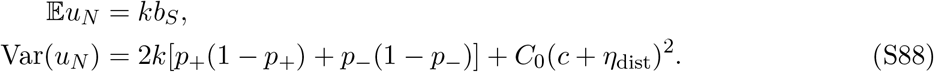

It can therefore quiet the novel population and improve specificity even while reduced write headroom lowers sensitivity. The optimum reflects both effects, rather than assignment concentration alone.

Distal readout coupling does not occur in the stationary or collision equations, so

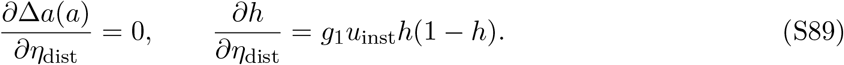

The memory and its matched no-write control have the same instructive-sign composition. Their distal translations cancel before the sigmoid,

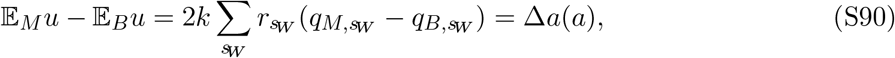

but the two plastic states occupy different parts of the nonlinear response. Consequently,

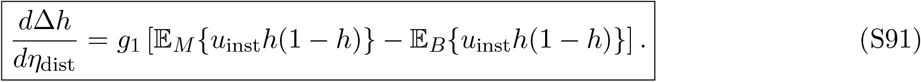

Distal input helps while the written memory unit carries more instructive-aligned mass on the responsive flank of the sigmoid; after the written response saturates, the matched no-write control can move faster and reduce the contrast. At the same time,

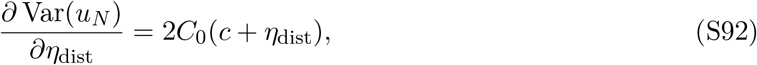

which is positive only on branches with *c* + *η*_dist_ *>* 0. In that regime, distal readout coupling expands the upper tail of novel responses and typically trades greater sensitivity against lower specificity. Between finite-lattice threshold crossings,

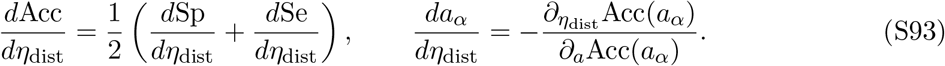

The memory-age-threshold derivative treats age as locally continuous and assumes a smooth, simple crossing. An interior accuracy optimum occurs when the marginal sensitivity gain balances the marginal specificity loss; direct evaluation is required at lattice or clipping discontinuities.

Overall, width mainly lowers the collision rate of each selected memory unit; input dimension and sparsity set finite-count reliability; *η*_bin_ and depression trade write strength against decay; *g*_2_ acts through the saturating selected-score moments; and *g*_1_, *b*_1_, Θ place the resulting memory-unit response laws relative to the decision boundary.

### 4.5 Hard-selection dual-depression Hebbian-control extension

The Hebbian control admits a parallel single-write reduced theory, but its interference geometry is different. It has no instructive matrix: the plastic drive controls both credit assignment and recognition,

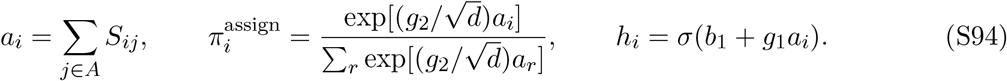

This plastic-state-dependent credit assignment echoes the self-overlap gate of Supplementary Note 1 but is not the same construction: the full Hebbian control also includes sparse inputs, competition among memory units, familiarity-dependent plasticity suppression and a distinct balanced depression rule. The extension retains exact-*k* inputs, single-level binary synapses, disabled familiarity-dependent plasticity suppression, one tagged write and read-only probes. It is therefore a matched stationary experiment, not a re-expression of the single-repeat capacity protocol with familiarity-dependent plasticity suppression.

#### Hard selection and balanced depression

Attainable plastic scores differ in steps of two. A sufficient width-aware condition for nonmaximal softmax mass to be negligible is

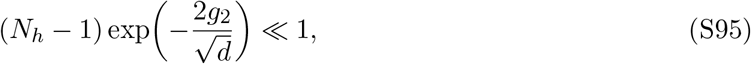

equivalently 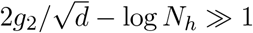 at large width. The softmax is then effectively a hard maximum. The theory chooses the selected memory unit from those with maximal *a*_*i*_, resolving an exact integer-score tie uniformly among the maximizers. This tie-breaking rule introduces neither an additional state nor a fitted parameter. The full model shares credit-assignment weight among tied selected memory units; categorical tie breaking preserves the mean exchangeability but removes those same-presentation cross-row correlations. The resulting disjoint selected- and unselected-memory-unit supports are therefore a property of the reduced theory.

With potentiation probability *η*_bin_, the two depression probabilities are fixed by

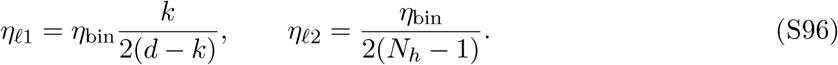

The selected memory unit receives LTD_1_ on inactive input coordinates and LTP on active input coordinates; every unselected memory unit receives LTD_2_ on active input coordinates. The full model’s within-presentation plasticity-pass order is LTD_1_ → LTD_2_ → LTP. In the unique-selection limit, the LTP and LTD_2_ supports are disjoint, so no same-coordinate repair term is needed. The expected attempted event masses are

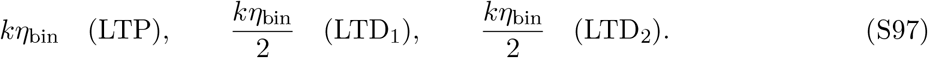

#### One-coordinate stationary solver

Without instructive-sign classes, the exchangeable product background has one unknown,

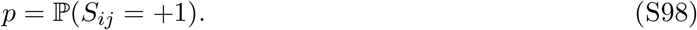

For the generic positive active count *J* of one memory unit, define

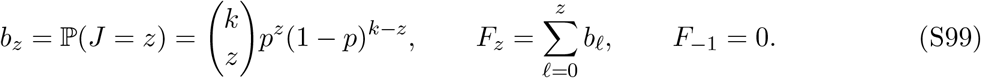

Conditional on *J* = *z*, the expected indicator that the memory unit is the selected memory unit after uniform maximum-tie breaking is

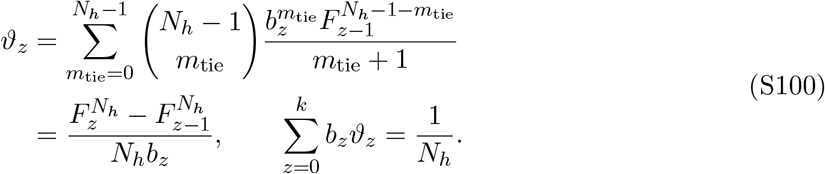

The first line enumerates the number of other rows that tie the focal row; the second is the corresponding order-statistic identity.

For an active coordinate with current plastic sign *s*_*S*_ ∈ {−1, +1}, its conditional selection probability is

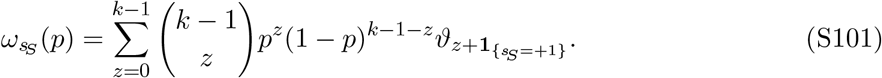

An inactive input coordinate is absent from the row score, so its memory unit is the selected memory unit for the pattern with probability 1*/N*_*h*_. The global per-presentation transition probabilities of one synapse are therefore

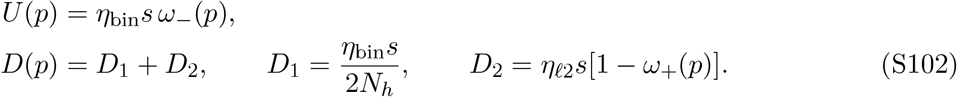

Stationary current balance gives the scalar equation

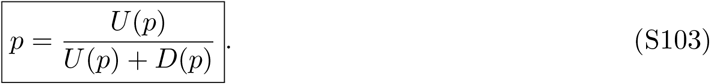

An equivalent order-statistic form makes uniqueness and computation transparent. If 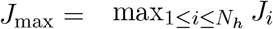 and *µ*_max_(*p*) = E*J*_max_, then

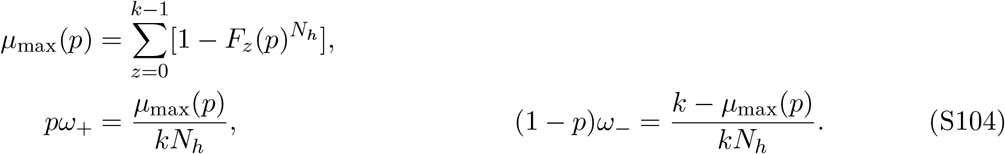

Substitution in equation (S103) gives

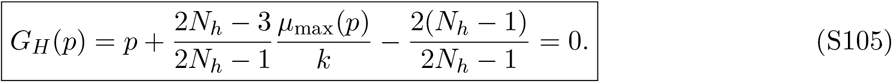

Moreover,

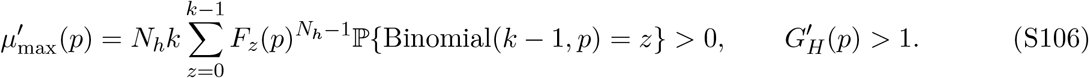

Hence there is one root in 0 *< p <* 1*/*2, obtainable with an *O*(*k*) safeguarded Newton solve. For nonzero plasticity the common factor *η*_bin_*s* cancels, so this stationary root depends only on *k* and *N*_*h*_; at *η*_bin_ = 0, all transition probabilities vanish and the dynamics do not select a stationary value. The role of *g*_2_ is only to establish the hard-selection condition in equation (S95). Exchangeability gives mean selection probability 1*/N*_*h*_ across the product ensemble, but does not assert that one fixed network realization uses every row equally over an indefinitely long stream; persistent row specialization is quantified directly in Fig. S2.

#### Selected- and unselected-memory-unit causal contrasts over memory age

Selection enriches the tagged coordinates before plasticity. Their positive fractions in the selected memory unit and in a specified unselected memory unit are

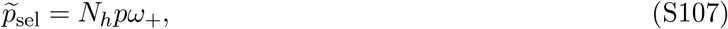

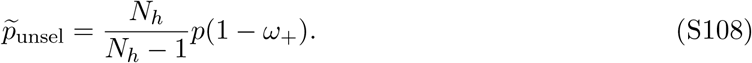

The one-shot memory and matched no-write states at immediate readout are

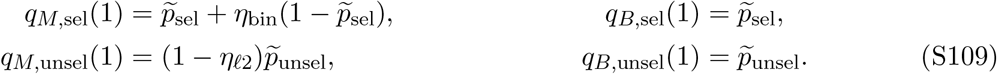

The one-shot write produces a positive causal engram contrast in the selected memory unit, while LTD_2_ produces a smaller negative causal contrast in each unselected memory unit. Every later presentation can modify every tagged memory unit, whether or not that memory unit is selected. Thus *U* and *D* already measure global per-presentation transition probabilities; multiplying them by another collision-rate factor would double-count sparse selection. Define

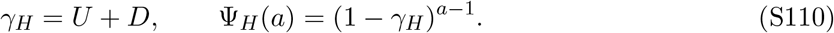

For *X* ∈ {*M, B*} and *ι* ∈ {sel, unsel},

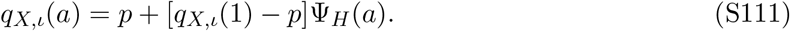

The population-level causal engram contrast is

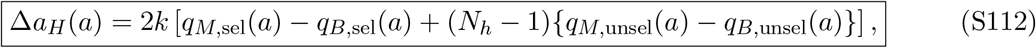

or, equivalently,

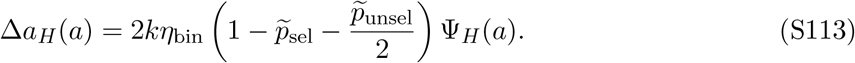

#### Finite-count readout

The maximum-conditioned count of the selected memory unit is narrower than a binomial law with the same mean, so its exact pre-write marginal is retained. The selected- and specified-unselected-memory-unit laws are

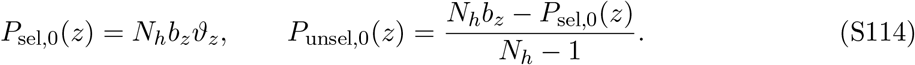

For the memory copy, each negative synapse in the selected memory unit potentiates independently with probability *η*_bin_, whereas each positive synapse in an unselected memory unit survives LTD_2_ with probability 1 − *η*_*ℓ*2_. Applying these binomial maps to *P*_sel,0_ and *P*_unsel,0_ defines the four immediate laws *P*_*M*,sel_(*z*; 1), *P*_*M*,unsel_(*z*; 1), *P*_*B*,sel_(*z*; 1) and *P*_*B*,unsel_(*z*; 1). Conditional on *z*_0_ positive synapses immediately after the tagged write, the stationary two-state kernel propagates the count to memory age *a* as

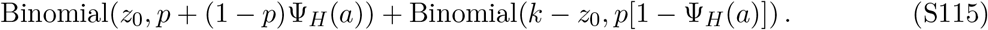

This finite law assumes conditionally independent coordinate transitions; exact-*k* inputs and common selection generate weak dependencies that are not retained. A novel memory unit remains distributed as *b*_*z*_, and each count maps to

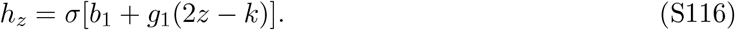

Denote the propagated laws by *P*_*X,ι*_(*z*; *a*), for *X* ∈ {*M, B*} and *ι* ∈ {sel, unsel}, and define 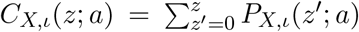 . In the general soft-soma calculation, a novel population keeps one generic *b*_*z*_ memory unit exactly and gamma-matches the other *N*_*h*_ − 1 generic memory units. A familiar population keeps the selected-memory-unit law *P*_*M*,sel_ exactly and separately gamma-matches the *N*_*h*_ − 1 unselected-memory-unit laws *P*_*M*,unsel_. These hypothesis-specific remainders are part of the population-independence closure. The matched no-write control laws *P*_*B*,sel_ and *P*_*B*,unsel_ define the causal engram contrast, but do not replace the generic novel population in this Hebbian readout.

A simpler threshold expression applies when the memory-unit response is increasing and *z*_⋆_ = min{*z* : *h*_*z*_ *>* Θ} exists with 1 ≤ *z*_⋆_ ≤ *k*. If

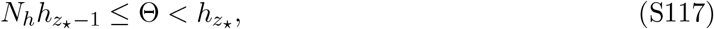

the population is familiar if and only if at least one memory unit reaches *z*_⋆_. The product closure gives

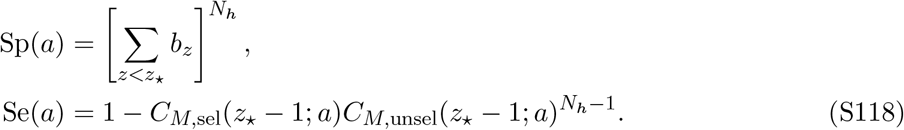

The single-trigger equivalence is exact under equation (S117); the product probability remains a population-independence closure because selection conditioning and shared future inputs correlate unselected memory units. Balanced accuracy and the memory-age threshold at the fixed readout threshold then follow from equations (S78) and (S79). Cases with no such *z*_⋆_, or with *z*_⋆_ = 0, are handled directly from the finite population law rather than by the single-trigger formula.

Within the homogeneous row manifold,

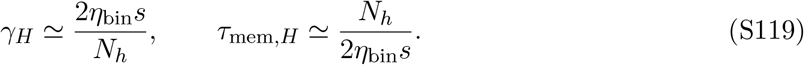

Width therefore extends retention approximately linearly, while *η*_bin_ trades initial write strength against decay. At fixed coding fraction, *d* acts mainly through finite-count reliability. Once the hard-selection condition holds, larger *g*_2_ does not change this reduction; *g*_1_, *b*_1_ and −*b*_2_*/w*_2_ instead determine how the count lattice is converted into a decision.

**Figure S2:**
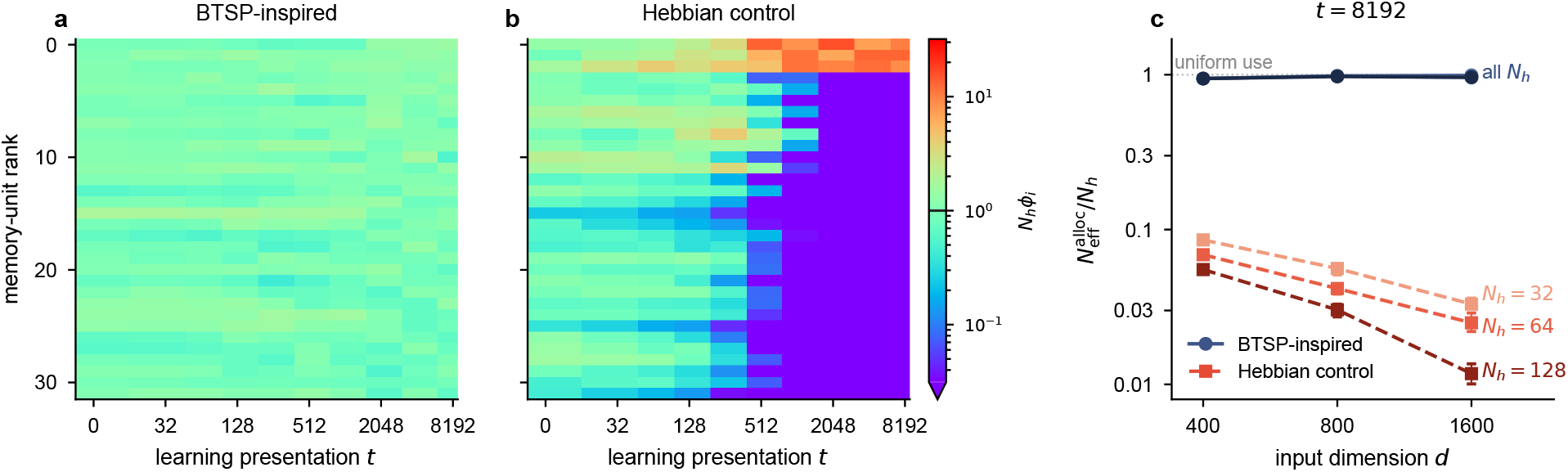
assignment-opportunity concentration in the rate-and-depth-matched networks. **a**,**b**, Relative assignment-opportunity share *N*_*h*_*ϕ*_*i*_ of every memory-unit row (equation (6); feedback-weighted credit-assignment mass on 2,048 fixed novel probes, evaluated at fixed network snapshots along one example continual stream) for the BTSP-inspired network (**a**) and the Hebbian control (**b**) at *d* = 400, *N*_*h*_ = 32 and *R* = 169. Rows are ranked by their late assignment-opportunity share; *N*_*h*_*ϕ*_*i*_ = 1 (black mark on the color bar) is uniform use, and the base-two logarithmic time axis extends to 8,192 presentations. **c**, Effective assigned-memory-unit fraction 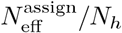 (equation (7)) at presentation 8,192 across the nine (*d, N*_*h*_) configurations of Fig. 4f. Navy, BTSP-inspired network; red, Hebbian control; shades, *N*_*h*_ = 32, 64 and 128 from light to dark; whiskers, 95% crossed-factor bootstrap intervals over 128 trajectories; dotted line, uniform use. In **a**,**b**, the Hebbian control is the rate-and-depth-matched Hebbian trace control of Fig. 5d; in **c**, the BTSP-inspired network uses each configuration’s saved *R*_0.90_ record, the Hebbian control uses its own *R*_0.90_ readout and gain parameters with *η*_bin_ and cascade depth matched to the BTSP-inspired network, and both networks see its *R*_0.90_ stream.

**Figure S3:**
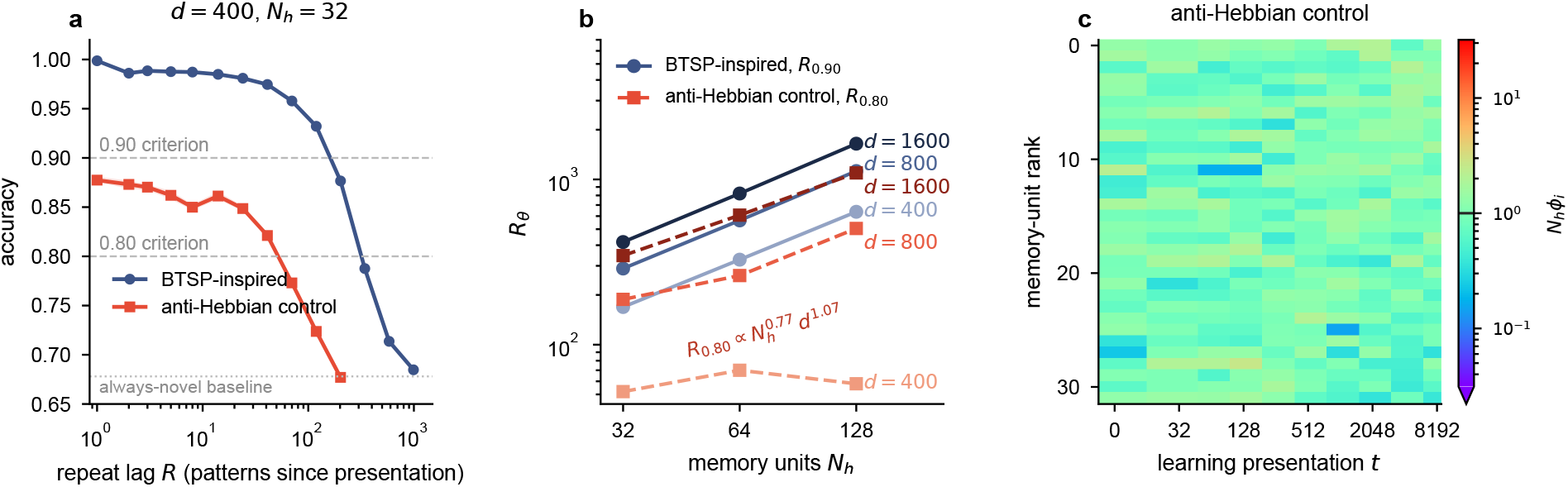
Anti-Hebbian plasticity alleviates assignment-opportunity concentration but reduces accuracy and capacity. **a**, Held-out protocol-weighted accuracy versus repeat lag for the BTSP-inspired network and anti-Hebbian control (*d* = 400, *N*_*h*_ = 32); dashed lines, 0.90 and 0.80 criteria; dotted line, always-novel baseline. The anti-Hebbian control remains below 0.90. **b**, Lag capacity versus memory-unit number and input dimension. Anti-Hebbian *R*_0.80_ scales with both dimensions similarly to the BTSP-inspired network’s *R*_0.90_ but remains substantially lower; the anti-Hebbian fit uses *d* = 800 and 1600. **c**, Relative assignment-opportunity share *N*_*h*_*ϕ*_*i*_ across memory-unit rows for the representative *R* = 169 stream of Supplementary Fig. S2, using the anti-Hebbian control optimized at its own *R*_0.80_. Depressing the selected memory unit redistributes subsequent writes and maintains broad memory-unit use. The anti-Hebbian control reverses the Hebbian event polarity (*η*_bin_ *<* 0); all other protocol details match Fig. 4.

## Supplementary Note 5: unified symbol dictionary

**Table S1:**
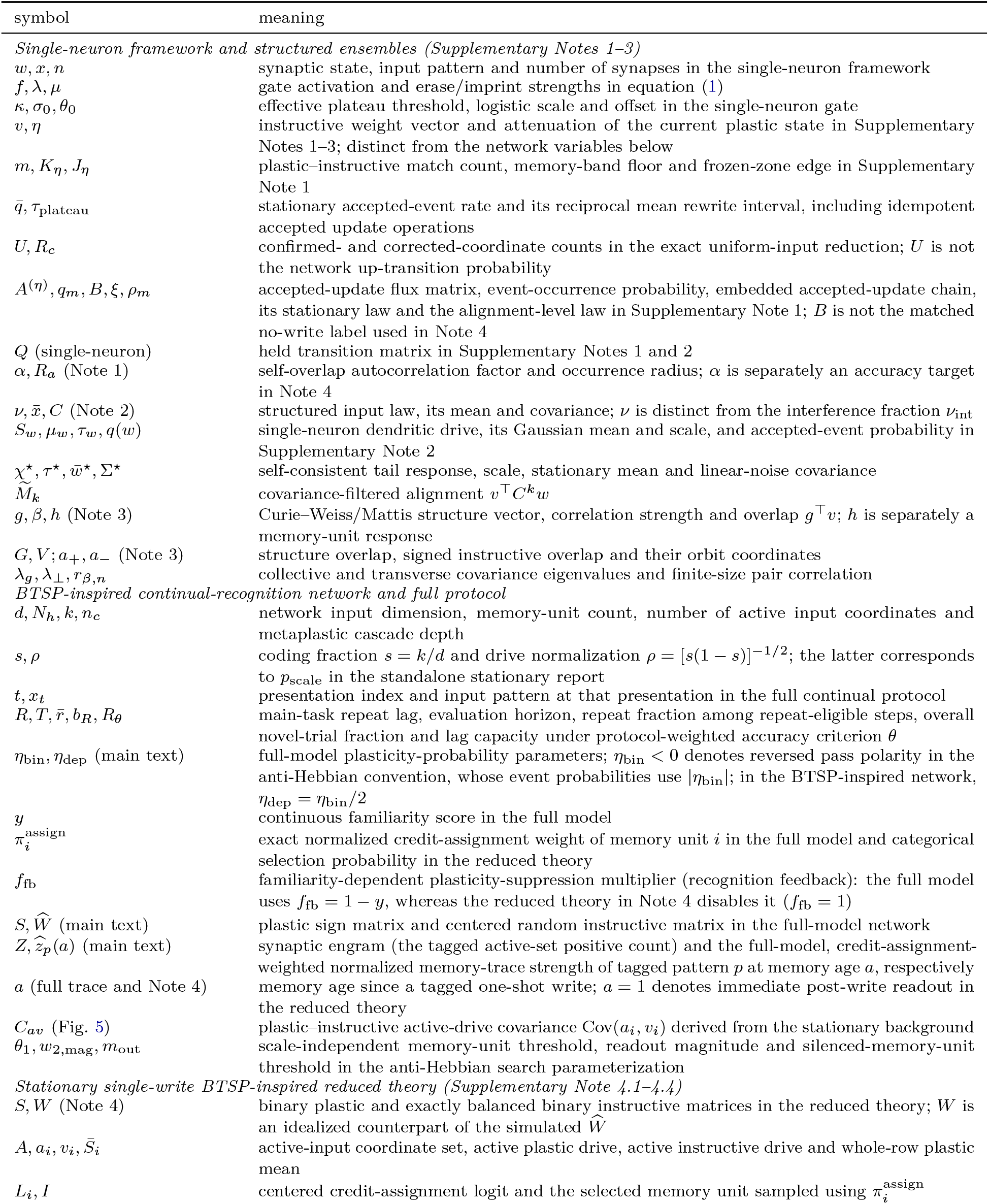
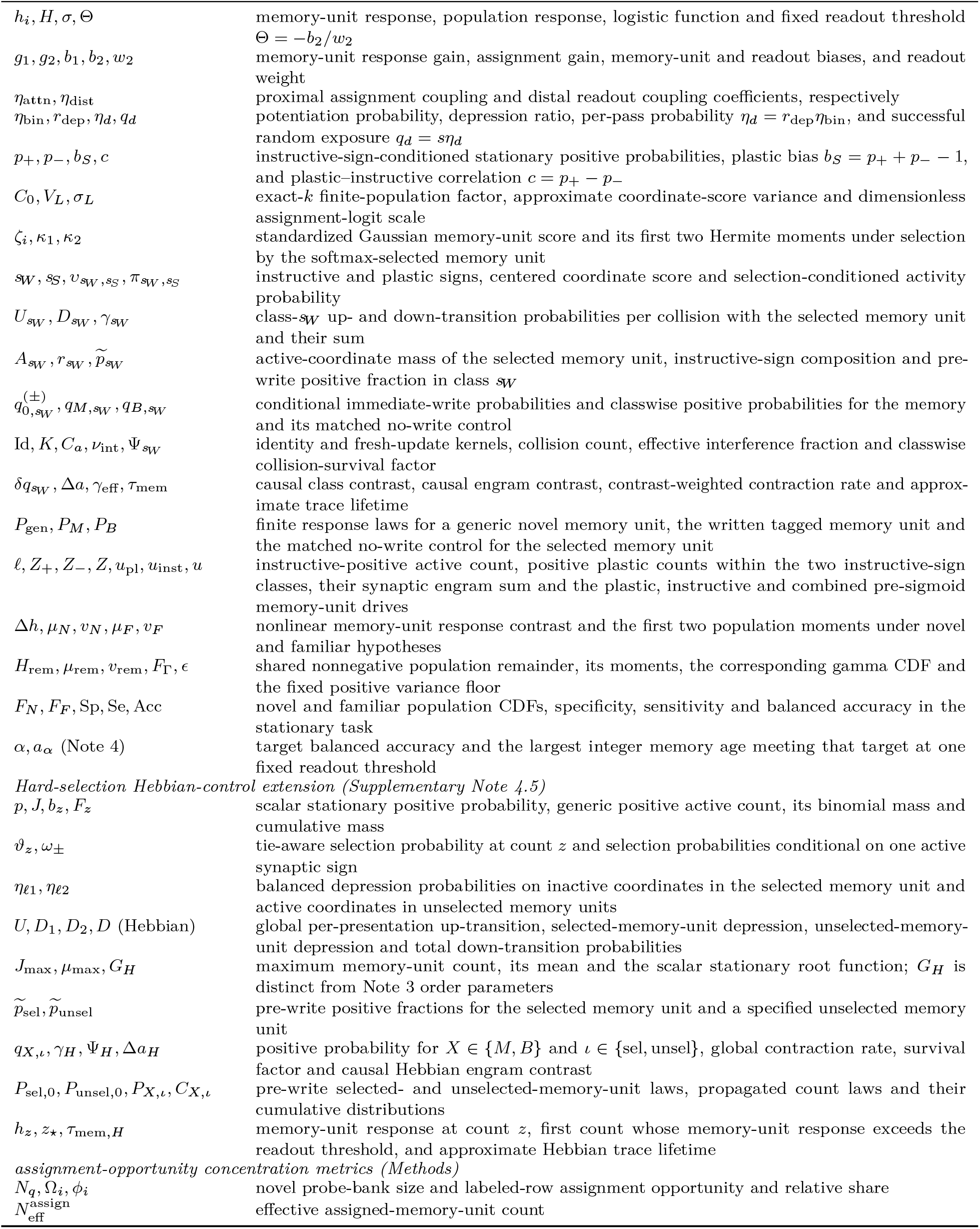
Principal notation used in the main text and Supplementary Notes. Reused symbols are qualified by mathematical scope; full continual-model quantities and reduced-theory quantities in Supplementary Note 4 are distinguished below.

## Notes

### Competing Interest Statement

The authors have declared no competing interest.

## References

[1] David Marr. Simple memory: a theory for archicortex. Philosophical Transactions of the Royal Society of London. Series B, Biological Sciences, 262(841):23–81, 1971. doi: 10.1098/rstb.1971.0078.

[2] Katie C. Bittner, Christine Grienberger, Sachin P. Vaidya, Aaron D. Milstein, John J. Macklin, Junghyup Suh, Susumu Tonegawa, and Jeffrey C. Magee. Conjunctive input processing drives feature selectivity in hippocampal CA1 neurons. Nature Neuroscience, 18:1133–1142, 2015.

[3] Katie C. Bittner, Aaron D. Milstein, Christine Grienberger, Sandro Romani, and Jeffrey C. Magee. Behavioral time scale synaptic plasticity underlies CA1 place fields. Science, 357:1033–1036, 2017.

[4] Jeffrey C. Magee. Behavioral timescale synaptic plasticity: properties, elements and functions. Nature Neuroscience, 29:520–534, 2026. doi: 10.1038/s41593-026-02214-2.

[5] Lionel Standing. Learning 10,000 pictures. Quarterly Journal of Experimental Psychology, 25:207–222, 1973.

[6] Timothy F. Brady, Talia Konkle, George A. Alvarez, and Aude Oliva. Visual long-term memory has a massive storage capacity for object details. Proceedings of the National Academy of Sciences USA, 105: 14325–14329, 2008.

[7] Michael McCloskey and Neal J. Cohen. Catastrophic interference in connectionist networks: the sequential learning problem. In Psychology of Learning and Motivation, volume 24, pages 109–165. Academic Press, 1989.

[8] Robert M. French. Catastrophic forgetting in connectionist networks. Trends in Cognitive Sciences, 3: 128–135, 1999.

[9] German I. Parisi, Ronald Kemker, Jose L. Part, Christopher Kanan, and Stefan Wermter. Continual lifelong learning with neural networks: a review. Neural Networks, 113:54–71, 2019.

[10] James Kirkpatrick, Razvan Pascanu, Neil Rabinowitz, Joel Veness, Guillaume Desjardins, Andrei A. Rusu, Kieran Milan, John Quan, Tiago Ramalho, Agnieszka Grabska-Barwinska, Demis Hassabis, Claudia Clopath, Dharshan Kumaran, and Raia Hadsell. Overcoming catastrophic forgetting in neural networks. Proceedings of the National Academy of Sciences USA, 114:3521–3526, 2017.

[11] Malcolm W. Brown and John P. Aggleton. Recognition memory: what are the roles of the perirhinal cortex and hippocampus? Nature Reviews Neuroscience, 2:51–61, 2001.

[12] Rafał Bogacz Malcolm W. Brown, and Christophe Giraud-Carrier. Model of familiarity discrimination in the perirhinal cortex. Journal of Computational Neuroscience, 10(1):5–23, 2001. doi: 10.1023/A:1008925909305.

[13] Rafał Bogacz and Malcolm W. Brown. Comparison of computational models of familiarity discrimination in the perirhinal cortex. Hippocampus, 13(4):494–524, 2003. doi: 10.1002/hipo.10093.

[14] João Sacramento and Andreas Wichert. Binary Willshaw learning yields high synaptic capacity for long-term familiarity memory. Biological Cybernetics, 106(2):123–133, 2012. doi: 10.1007/s00422-012-0488-4.

[15] Danil Tyulmankov, Guangyu Robert Yang, and L. F. Abbott. Meta-learning synaptic plasticity and memory addressing for continual familiarity detection. Neuron, 110:544–557, 2022.

[16] Jin-Hee Han, Steven A. Kushner, Adelaide P. Yiu, Christy J. Cole, Anna Matynia, Robert A. Brown, Rachael L. Neve, John F. Guzowski, Alcino J. Silva, and Sheena A. Josselyn. Neuronal competition and selection during memory formation. Science, 316(5823):457–460, 2007. doi: 10.1126/science.1139438.

[17] Alcino J. Silva, Yu Zhou, Thomas Rogerson, Justin Shobe, and J. Balaji. Molecular and cellular approaches to memory allocation in neural circuits. Science, 326(5951):391–395, 2009. doi: 10.1126/science.1174519.

[18] Jeffrey C. Magee and Christine Grienberger. Synaptic plasticity forms and functions. Annual Review of Neuroscience, 43:95–117, 2020.

[19] Aaron D. Milstein, Yiding Li, Katie C. Bittner, Christine Grienberger, Ivan Soltesz, Jeffrey C. Magee, and Sandro Romani. Bidirectional synaptic plasticity rapidly modifies hippocampal representations. eLife, 10:e73046, 2021.

[20] Yiding Li, John J. Briguglio, Sandro Romani, and Jeffrey C. Magee. Mechanisms of memory-supporting neuronal dynamics in hippocampal area CA3. Cell, 187(24):6804–6819.e21, 2024. doi: 10.1016/j.cell.2024.09.041.

[21] Hiroto Takahashi and Jeffrey C. Magee. Pathway interactions and synaptic plasticity in the dendritic tuft regions of CA1 pyramidal neurons. Neuron, 62:102–111, 2009.

[22] Christine Grienberger and Jeffrey C. Magee. Entorhinal cortex directs learning-related changes in CA1 representations. Nature, 611:554–562, 2022.

[23] Wulfram Gerstner, Marco Lehmann, Vasiliki Liakoni, Dane Corneil, and Johanni Brea. Eligibility traces and plasticity on behavioral time scales: experimental support of neoHebbian three-factor learning rules. Frontiers in Neural Circuits, 12:53, 2018. doi: 10.3389/fncir.2018.00053.

[24] Donald O. Hebb. The Organization of Behavior. Wiley, New York, 1949.

[25] John J. Hopfield. Neural networks and physical systems with emergent collective computational abilities. Proceedings of the National Academy of Sciences USA, 79:2554–2558, 1982.

[26] Daniel J. Amit, Hanoch Gutfreund, and Haim Sompolinsky. Storing infinite numbers of patterns in a spin-glass model of neural networks. Physical Review Letters, 55:1530–1533, 1985.

[27] Daniel J. Amit, Hanoch Gutfreund, and Haim Sompolinsky. Statistical mechanics of neural networks near saturation. Annals of Physics, 173(1):30–67, 1987. doi: 10.1016/0003-4916(87)90092-3.

[28] David J. Willshaw, O. Peter Buneman, and H. Christopher Longuet-Higgins. Non-holographic associative memory. Nature, 222:960–962, 1969. doi: 10.1038/222960a0.

[29] M. V. Tsodyks and M. V. Feigel’man. The enhanced storage capacity in neural networks with low activity level. Europhysics Letters, 6(2):101–105, 1988. doi: 10.1209/0295-5075/6/2/002.

[30] Daniel J. Amit, Hanoch Gutfreund, and Haim Sompolinsky. Information storage in neural networks with low levels of activity. Physical Review A, 35(5):2293–2303, 1987. doi: 10.1103/PhysRevA.35.2293.

[31] David Golomb, Nava Rubin, and Haim Sompolinsky. Willshaw model: associative memory with sparse coding and low firing rates. Physical Review A, 41(4):1843–1854, 1990. doi: 10.1103/PhysRevA.41.1843.

[32] Jean-Pierre Nadal, Gérard Toulouse, Jean-Pierre Changeux, and Stanislas Dehaene. Networks of formal neurons and memory palimpsests. Europhysics Letters, 1(10):535–542, 1986. doi: 10.1209/0295-5075/1/10/008.

[33] Giorgio Parisi. A memory which forgets. Journal of Physics A: Mathematical and General, 19:L617–L620, 1986.

[34] M. V. Tsodyks. Associative memory in neural networks with binary synapses. Modern Physics Letters B, 4(11):713–716, 1990. doi: 10.1142/S0217984990000891.

[35] Daniel J. Amit and Stefano Fusi. Constraints on learning in dynamic synapses. Network: Computation in Neural Systems, 3(4):443–464, 1992. doi: 10.1088/0954-898X/3/4/008.

[36] Daniel J. Amit and Stefano Fusi. Learning in neural networks with material synapses. Neural Computation, 6:957–982, 1994.

[37] Stefano Fusi and Larry F. Abbott. Limits on the memory storage capacity of bounded synapses. Nature Neuroscience, 10:485–493, 2007.

[38] Adam B. Barrett and Mark C. W. van Rossum. Optimal learning rules for discrete synapses. PLoS Computational Biology, 4(11):e1000230, 2008. doi: 10.1371/journal.pcbi.1000230.

[39] Stefano Fusi, Patrick J. Drew, and Larry F. Abbott. Cascade models of synaptically stored memories. Neuron, 45:599–611, 2005.

[40] Marcus K. Benna and Stefano Fusi. Computational principles of synaptic memory consolidation. Nature Neuroscience, 19:1697–1706, 2016.

[41] Subhaneil Lahiri and Surya Ganguli. A memory frontier for complex synapses. In Advances in Neural Information Processing Systems, volume 26, pages 1034–1042, 2013.

[42] Walter Senn and Stefano Fusi. Learning only when necessary: better memories of correlated patterns in networks with bounded synapses. Neural Computation, 17(10):2106–2138, 2005. doi: 10.1162/0899766054615644.

[43] Ian Cone and Harel Z. Shouval. Behavioral time scale plasticity of place fields: mathematical analysis. Frontiers in Computational Neuroscience, 15:640235, 2021.

[44] Yujie Wu and Wolfgang Maass. A simple model for behavioral time scale synaptic plasticity (BTSP) pro-vides content addressable memory with binary synapses and one-shot learning. Nature Communications, 16:342, 2025.

[45] Xundong E. Wu and Bartlett W. Mel. Capacity-enhancing synaptic learning rules in a medial temporal lobe online learning model. Neuron, 62:31–41, 2009.

[46] Aaron D. Milstein, Erik B. Bloss, Pierre F. Apostolides, Sachin P. Vaidya, Geoffrey A. Dilly, Boris V. Zemelman, and Jeffrey C. Magee. Inhibitory gating of input comparison in the CA1 microcircuit. Neuron, 87(6):1274–1289, 2015. doi: 10.1016/j.neuron.2015.08.025.

[47] Evan P. Campbell, Lisandro Martin, Jeffrey C. Magee, and Christine Grienberger. Learning-dependent feedback by OLM interneurons shapes CA1 representations. bioRxiv, 2026. doi: 10.64898/2025.12.21.695825. Preprint, version 2.

[48] Yaniv Ziv, Laurie D. Burns, Eric D. Cocker, Elizabeth O. Hamel, Kunal K. Ghosh, Lacey J. Kitch, Abbas El Gamal, and Mark J. Schnitzer. Long-term dynamics of CA1 hippocampal place codes. Nature Neuroscience, 16:264–266, 2013.

[49] Michael E. Rule, Timothy O’Leary, and Christopher D. Harvey. Causes and consequences of representational drift. Current Opinion in Neurobiology, 58:141–147, 2019.

[50] Antoine D. Madar, Anqi Jiang, Can Dong, and Mark E. J. Sheffield. Synaptic plasticity rules driving representational shifting in the hippocampus. Nature Neuroscience, 28:848–860, 2025. doi: 10.1038/s41593-025-01894-6.

[51] Sachin P. Vaidya, Guanchun Li, Raymond A. Chitwood, Yiding Li, and Jeffrey C. Magee. Formation of an expanding memory representation in the hippocampus. Nature Neuroscience, 28:1510–1518, 2025. doi: 10.1038/s41593-025-01986-3.

[52] Rémi Monasson. Properties of neural networks storing spatially correlated patterns. Journal of Physics A: Mathematical and General, 25(13):3701–3720, 1992. doi: 10.1088/0305-4470/25/13/019.

[53] Barak Blumenfeld, Son Preminger, Dov Sagi, and Misha Tsodyks. Dynamics of memory representations in networks with novelty-facilitated synaptic plasticity. Neuron, 52(2):383–394, 2006. doi: 10.1016/j.neuron.2006.08.016.

[54] Daniel C. Mattis. Solvable spin systems with random interactions. Physics Letters A, 56:421–422, 1976.

[55] Iain M. Johnstone. On the distribution of the largest eigenvalue in principal components analysis. Annals of Statistics, 29:295–327, 2001.

[56] Jinho Baik, Gérard Ben Arous, and Sandrine Péché. Phase transition of the largest eigenvalue for nonnull complex sample covariance matrices. Annals of Probability, 33:1643–1697, 2005.

[57] Christoph von der Malsburg. Self-organization of orientation sensitive cells in the striate cortex. Kybernetik, 14:85–100, 1973.

[58] David E. Rumelhart and David Zipser. Feature discovery by competitive learning. Cognitive Science, 9: 75–112, 1985.

[59] Sheena A. Josselyn and Susumu Tonegawa. Memory engrams: recalling the past and imagining the future. Science, 367(6473):eaaw4325, 2020. doi: 10.1126/science.aaw4325.

[60] Rafał Bogacz and Malcolm W. Brown. An anti-Hebbian model of familiarity discrimination in the perirhinal cortex. Neurocomputing, 52–54:1–6, 2003. doi: 10.1016/S0925-2312(02)00738-5.

[61] Andrea Greve, David C. Sterratt, David I. Donaldson, David J. Willshaw, and Mark C. W. van Rossum. Optimal learning rules for familiarity detection. Biological Cybernetics, 100(1):11–19, 2009. doi: 10.1007/s00422-008-0275-4.

[62] Nimrod Shaham, Jay Chandra, Gabriel Kreiman, and Haim Sompolinsky. Stochastic consolidation of lifelong memory. Scientific Reports, 12:13107, 2022. doi: 10.1038/s41598-022-16407-9.

[63] Pentti Kanerva. Sparse Distributed Memory. MIT Press, Cambridge, MA, 1988. ISBN 978-0-262-11132-4.

[64] Lee Susman, Naama Brenner, and Omri Barak. Stable memory with unstable synapses. Nature Communications, 10:4441, 2019. doi: 10.1038/s41467-019-12306-2.

[65] Stefano Fusi and Walter Senn. Eluding oblivion with smart stochastic selection of synaptic updates. Chaos, 16(2):026112, 2006. doi: 10.1063/1.2213587.

[66] Friedemann Zenke, Ben Poole, and Surya Ganguli. Continual learning through synaptic intelligence. In Proceedings of the 34th International Conference on Machine Learning, volume 70 of PMLR, pages 3987–3995, 2017.

[67] David J. Willshaw and Peter Dayan. Optimal plasticity from matrix memories: what goes up must come down. Neural Computation, 2(1):85–93, 1990. doi: 10.1162/neco.1990.2.1.85.

[68] Takuya Akiba, Shotaro Sano, Toshihiko Yanase, Takeru Ohta, and Masanori Koyama. Optuna: a next-generation hyperparameter optimization framework. In Proceedings of the 25th ACM SIGKDD International Conference on Knowledge Discovery and Data Mining, pages 2623–2631, 2019.

[69] John G. Kemeny and J. Laurie Snell. Finite Markov Chains. Springer, New York, 1976.

